# An integrated view of baseline protein expression in human tissues

**DOI:** 10.1101/2021.09.10.459811

**Authors:** Ananth Prakash, David García-Seisdedos, Shengbo Wang, Deepti Jaiswal Kundu, Andrew Collins, Nancy George, Pablo Moreno, Irene Papatheodorou, Andrew R. Jones, Juan Antonio Vizcaíno

**Affiliations:** European Molecular Biology Laboratory - European Bioinformatics Institute (EMBL-EBI), Wellcome Genome Campus, Hinxton, Cambridge, CB10 1SD. United Kingdom; Open Targets, Wellcome Genome Campus, Hinxton, Cambridge, CB10 1SD. United Kingdom; Institute of Systems, Molecular and Integrative Biology, University of Liverpool, Liverpool L69 7ZB, United Kingdom

**Author notes:** Corresponding authors: Dr. Ananth Prakash. European Molecular Biology Laboratory, European Bioinformatics Institute (EMBL-EBI), Wellcome Trust Genome Campus, Hinxton, Cambridge, CB10 1SD, UK.; Prof. Andrew R. Jones. Institute of Systems, Molecular and Integrative Biology, University of Liverpool, Liverpool L69 7ZB, United Kingdom.; Dr. Juan Antonio Vizcaíno. European Molecular Biology Laboratory, European Bioinformatics Institute (EMBL-EBI), Wellcome Trust Genome Campus, Hinxton, Cambridge, CB10 1SD, UK.

**Keywords:** Mass spectrometry, quantitative proteomics, public data re-use, human proteome

## Abstract

The availability of proteomics datasets in the public domain, and in the PRIDE database in particular, has increased dramatically in recent years. This unprecedented large-scale availability of data provides an opportunity for combined analyses of datasets to get organism-wide protein abundance data in a consistent manner. We have reanalysed 24 public proteomics datasets from healthy human individuals, to assess baseline protein abundance in 31 organs. We defined tissue as a distinct functional or structural region within an organ. Overall, the aggregated dataset contains 67 healthy tissues, corresponding to 3,119 mass spectrometry runs covering 498 samples, coming from 489 individuals.

We compared protein abundances between the different organs and studied the distribution of proteins across organs. We also compared the results with data generated in analogous studies. We also performed gene ontology and pathway enrichment analyses to identify organ-specific enriched biological processes and pathways. As a key point, we have integrated the protein abundance results into the resource Expression Atlas, where it can be accessed and visualised either individually or together with gene expression data coming from transcriptomics datasets. We believe this is a good mechanism to make proteomics data more accessible for life scientists.

## Introduction

High-throughput mass spectrometry (MS)-based proteomics approaches have matured and generalised significantly, becoming an essential tool in biological research, sometimes together with other “omics” approaches such as genomics and transcriptomics. It is now commonplace to make quantitative measurements of 2,000-3,000 proteins in a single LC-MS run, and typically 6,000-7,000 proteins in workflows with fractionation. The most used experimental approach is Data Dependent Acquisition (DDA) bottom-up proteomics. Among existing DDA quantitative proteomics approaches, label-free is very popular, although labelled-approaches such as metabolic-labelling (e.g., SILAC) and especially techniques based on the isotopic labelling of peptides (e.g., TMT) are growing in importance. In bottom-up experiments, proteins are first digested into peptides using an enzyme (e.g., trypsin), and typically several peptides are required per protein to give confidence in the measurement of protein-level quantification across samples. Measured peptide intensity is correlated with absolute protein abundance, but there can be differences depending on individual peptides due to the considerable variation in the ionisation efficiency of these peptides. Different peptides can also be detected in different studies, giving rise to variability in protein abundance. One further challenge in quantitative proteomics relates to the “protein inference” problem [1]. In brief, many peptide sequences cannot be uniquely mapped to a single protein due to common conserved sequences present in different gene families (paralogs). During the last decade technological advances in MS have led to a large number of studies that have analysed protein abundances across various human tissues and organs [2–5]. These efforts are complemented by the comprehensive characterisation of the human proteome performed within the Human Proteome Project (HPP) [6–8], although the HPP has been focused on the identification of proteins, without performing any quantitative analysis.

In parallel with the technical developments in chromatography, MS and bioinformatics, the proteomics community has evolved to largely support open data practices. In brief, this means that datasets are released alongside publications, allowing other groups to check findings or re-analyse data with different approaches to generate new findings. Therefore, in recent years, the amount and variety of shared datasets in the public domain has grown dramatically. This was driven by the establishment and maturation of reliable proteomics data repositories, in tandem with policy recommendations by scientific journals and funding agencies.

The PRIDE database [9], which is one of the founding members of the global ProteomeXchange consortium [10], is currently the largest resource worldwide for public proteomics data deposition. As of October 2022, PRIDE hosts more than 29,500 datasets. Of those, human datasets are by far the majority, representing approximately 40% of all datasets. Public datasets stored in PRIDE (or in other resources) present an opportunity to be systematically reanalysed and integrated, in order to confirm the original results potentially in a more robust manner, obtain new insights, generate new hypotheses, and even be able to answer biologically relevant questions orthogonal to those posed in the original studies. Such integrative meta-analyses have already been successfully employed especially in genomics and transcriptomics [11–13]. Therefore, the large availability of public datasets has triggered different types of data re-use activities, including “big data” approaches (e.g. [14–16]) and the establishment of new data resources using re-analysed public datasets as the basis [17–19]. In this context of data re-use, the main interest of PRIDE is to disseminate and integrate proteomics data into popular added-value bioinformatics resources at the European Bioinformatics Institute (EMBL-EBI) such as Expression Atlas [20] (for quantitative proteomics data), Ensembl [21] (proteogenomics) and UniProt [7] (protein sequences information including post-translational modifications (PTMs)). The overall aim is to enable life scientists (including those who are non-experts in proteomics) to have improved access to proteomics-derived information. Expression Atlas (https://www.ebi.ac.uk/gxa/home) is an added-value resource that enables easy access to integrated information about gene and recently protein expression across species, tissues, cells, experimental conditions and diseases. The Expression Atlas ‘bulk’ Atlas has two sections: baseline and differential atlas. Protein abundance results derived from the reanalysis of DDA public datasets of different sources have started to be incorporated into Expression Atlas. The availability of such results in Expression Atlas makes proteomics abundance data integrated with transcriptomics information in the web interface. We have performed two DDA studies of this type so far. First of all, we reported the reanalysis and integration into Expression Atlas of 11 public quantitative datasets coming from cell lines and human tumour samples [22]. Additionally, we have recently reported the reanalysis and integration of 23 datasets coming from mouse and rat tissues in baseline conditions [23].

There are other public resources providing access to reanalysed MS-based quantitative proteomics datasets. ProteomicsDB [24] provides access to human protein abundance data in addition to other recent (multi-omic) studies carried out on model organisms. Many additional human datasets coming from human tissues have been made publicly available in recent years. Within the HPP, it is important to highlight that ProteomeXchange resources PeptideAtlas [25] and MassIVE provide peptide and protein identifications derived from the reanalysis of public human datasets, but their main focus is not quantitative data. Additionally, antibody-based protein abundance information can be accessed *via* the Human Protein Atlas (HPA) [4]. Here, we report the reanalysis and integration of 24 public human label-free datasets, and the incorporation of the results into Expression Atlas as baseline studies.

## Experimental Procedures

### Datasets

As of September 2020, 3,930 public MS human proteomics datasets were publicly available in PRIDE. We manually filtered these 3,930 human datasets to select suitable datasets for downstream analyses by applying several selection criteria. These selection criteria for the datasets to be reanalysed were: i) experimental data from healthy tissues in baseline conditions coming from label-free studies where no PTM-enrichment had been performed; ii) experiments performed on Thermo Fisher Scientific instruments (LTQ Orbitrap, LTQ Orbitrap Elite, LTQ Orbitrap Velos, LTQ Orbitrap XL ETD, LTQ-Orbitrap XL ETD, Orbitrap Fusion and Q-Exactive), because they represent the larger proportion of the relevant public datasets available, and we preferred to avoid the heterogeneity introduced by using data coming from different MS vendors; iii) availability of detailed sample metadata in the original publication, or after contacting the original submitters; and iv) our previous experience in the team working with some datasets, which were discarded because they were not considered to be usable (data not shown). As a result 16 human datasets from PRIDE (Table 1). Additionally, 8 datasets coming from human brain samples (also generated in Thermo Fisher Scientific instruments) were downloaded from a large Alzheimer’s Disease (AD) dataset described in [26], which was available via the AMP-AD Knowledge Portal (https://adknowledgeportal.synapse.org/). Due to ethical related issues, the AD datasets from the AMP-AD Knowledge Portal are available under a controlled access agreement (i.e., data made available only to approved users of the data included in the AMP-AD Knowledge Portal) and were downloaded after obtaining the required authorisation.

**Table 1.** List of proteomics datasets that were reanalysed. *Dataset identifiers starting with ‘PXD’ come from the PRIDE database and those identifiers starting by ‘syn’ come from the AMP-AD Knowledge Portal. ^§^Only normal samples within this dataset are reported in this study. However, results from both normal and disease samples are available in Expression Atlas. Unique protein sample batches available in any given dataset are considered as individual samples (for example, dataset E-PROT-34 (PXD004143) consists of four experiment batches, where materials from two donors are each digested with LysC and trypsin, and therefore these four unique batches are considered as four different samples). † Numbers after post-processing. The proteomics results in Expression Atlas can be accessed using the link: https://www.ebi.ac.uk/gxa/experiments/E-PROT-XX/Results, where XX should be replaced by the E-PROT accession number shown in the table. The raw proteomics datasets in PRIDE can be accessed using the link: https://www.ebi.ac.uk/pride/archive/projects/PXDxxxxxx, where PXDxxxxxx should be replaced by the PRIDE dataset identifier shown in the table.

| Expression Atlas accession number | Proteomics dataset identifier* | Tissues | Organs | Mass spectrometer | Number of MS runs | Number of samples | Fractionation | Number of protein groups† | Number of peptides† | Number of unique peptides† | Number of unique genes (canonical proteins) mapped† |
| --- | --- | --- | --- | --- | --- | --- | --- | --- | --- | --- | --- |
| E-PROT-29 | PXD010154 <sup>[36]</sup> | Adrenal gland, Bone marrow, Brain, Colon, Duodenum, Esophagus, Fallopian tube oviduct, Fat adipose tissue, Gallbladder, Heart, Kidney, Liver, Lung, Lymph node, Ovary, Pancreas, Pituitary hypophysis, Placenta, Prostate, Rectum, Salivary gland, Small intestine, Smooth muscle, Spleen, Stomach, Testis, Thyroid, Tonsil, Urinary bladder, Uterine endometrium, Vermiform appendix | Adrenal gland, Bone marrow, Brain, Colon, Duodenum, Esophagus, Fallopian tube oviduct, Fat adipose tissue, Gallbladder, Heart, Kidney, Liver, Lung, Lymph node, Ovary, Pancreas, Placenta, Prostate, Rectum, Salivary gland, Small intestine, Smooth muscle, Spleen, Stomach, Testis, Thyroid, Tonsil, Urinary bladder, Uterine endometrium, Vermiform appendix | Q-Exactive Plus | 1,795 | 37 | Yes | 19,382 | 874,825 | 346,894 | 12,622 |
| E-PROT-33 | PXD005819 <sup>[37]</sup> | Anterior pituitary gland (adenohypophysis) | Brain | LTQ-Orbitrap Velos | 33 | 2 | Yes | 1,936 | 12,112 | 10,174 | 1,207 |
| E-PROT-34 | PXD004143 <sup>[38]</sup> | Dorsolateral prefrontal cortex | Brain | Thermo Orbitrap Fusion | 80 | 4 | Yes | 9,929 | 203,798 | 124,355 | 7,633 |
| E-PROT-35 | PXD006233 <sup>[39]</sup> | Middle temporal lobe | Brain | Orbitrap Elite | 192 | 6 | Yes | 7,030 | 62,455 | 46,786 | 5,264 |
| E-PROT-36 | PXD012755 <sup>[40]</sup> | Cerebellar hemispheric cortex, Occipital cortex | Brain | OrbitrapVelos Elite | 15 | 15 | No | 3,033 | 25,514 | 20,125 | 2,264 |
| E-PROT-40 | PXD001608 <sup>[41]</sup> | Colon | Colon | Q-Exactive Plus | 30 | 10 | No | 5,943 | 73,053 | 56,756 | 4,789 |
| E-PROT-41 | PXD002029 <sup>[42]</sup> | Colon | Colon | Q-Exactive Plus | 24 | 8 | No | 4,608 | 25,300 | 22,032 | 3,211 |
| E-PROT-42 | PXD000547,<br>PXD000548 <sup>[43]</sup> | Corpus callosum, Anterior temporal lobe | Brain | LTQ-Orbitrap XL | 40, 40 | 2, 2 | Yes | 1,353, 1,873 | 8,250, 12,503 | 6,739, 10,071 | 868, 1,304 |
| E-PROT-43 | PXD010271 <sup>[44]</sup> | Liver, Ovary, Pancreas, Substantia nigra | Liver, Ovary, Pancreas, Brain | Velos Orbitrap, Q-Exactive | 55 | 54 | No | 5,927 | 75,320 | 50,652 | 4,528 |
| E-PROT-44 | PXD004332 <sup>[45]</sup> | Pineal gland | Brain | LTQ-Orbitrap Velos | 56 | 3 | Yes | 4,953 | 49,455 | 38,884 | 3,692 |
| E-PROT-45 | PXD006675 <sup>[46]</sup> | Aorta, Aortic valve, Atrial septum, Inferior vena cava, Left atrium, Left ventricle, Mitral valve, Pulmonary artery, Pulmonary valve, Pulmonary vein, Right atrium, Right ventricle, Tricuspid valve, Ventricular septum | Cardiovascular system (Heart) | Q-Exactive HF | 347 | 42 | Yes | 9,160 | 161,943 | 98,692 | 7,602 |
| E-PROT-46 § | PXD008934 <sup>[47]</sup> | Left ventricle | Cardiovascular system (Heart) | Q-Exactive | 7 | 7 | No | 2,977 | 31,755 | 25,704 | 2,294 |
| E-PROT-51 § | syn6038852 <sup>[48]</sup> | Dorsolateral prefrontal cortex | Brain | Q-Exactive Plus | 11 | 11 | No | 4,962 | 56,558 | 41,312 | 3,803 |
| E-PROT-52 <sup>§</sup> | syn21444980 <sup>[26]</sup> | Dorsolateral prefrontal cortex | Brain | Q-Exactive Plus | 83 | 83 | No | 3,846 | 55,290 | 39,470 | 3,127 |
| E-PROT-53 <sup>§</sup> | syn7204174 <sup>[49]</sup> | Dorsolateral prefrontal cortex | Brain | Q-Exactive Plus | 26 | 26 | No | 5,658 | 101,460 | 64,060 | 4,454 |
| E-PROT-54 <sup>§</sup> | syn3606087 <sup>[50]</sup> | Dorsolateral prefrontal cortex | Brain | Q-Exactive Plus | 11 | 11 | No | 4,751 | 58,302 | 40,469 | 3,626 |
| E-PROT-55 <sup>§</sup> | syn4624471 <sup>[50]</sup> | Precuneus | Brain | Q-Exactive Plus | 13 | 13 | No | 4,829 | 60,129 | 42,179 | 3,695 |
| E-PROT-56 <sup>§</sup> | syn7431984 <sup>[51]</sup> | Temporal cortex | Brain | Q-Exactive Plus | 31 | 31 | No | 5,997 | 113,820 | 68,951 | 4,675 |
| E-PROT-57 <sup>§</sup> | syn6038797 <sup>[52]</sup> | Frontal pole | Brain | Q-Exactive Plus | 53 | 53 | No | 5,812 | 91,285 | 59,701 | 4,420 |
| E-PROT-58 <sup>§</sup> | syn21443008 <sup>[26]</sup> | Dorsolateral prefrontal cortex | Brain | Q-Exactive Plus | 47 | 47 | No | 5,050 | 86,685 | 55,386 | 4,053 |
| E-PROT-61 <sup>§</sup> | PXD012131 <sup>[53]</sup> | Amygdala, Caudate nucleus, Cerebellum, Entorhinal cortex, Inferior parietal lobule, Middle frontal gyrus, Neocortex, Superior temporal gyrus, Thalamus, Visual cortex | Brain | Orbitrap Fusion Lumos | 114 | 15 | Yes | 10,097 | 192,397 | 113,310 | 7,826 |
| E-PROT-63 | PXD020187 <sup>[54]</sup> | Umbilical artery | Umbilical artery | nano-ESI Orbitrap-Elite | 10 | 10 | No | 1,505 | 10,365 | 7,702 | 933 |
| E-PROT-65 <sup>§</sup> | PXD015079 <sup>[55]</sup> | Prefrontal cortex, Vermiform appendix | Brain, Vermiform appendix | Q-Exactive HF-X | 6 | 6 | No | 2,313 | 24,162 | 18,892 | 1,694 |
| **TOTAL** | **24 datasets** | **67 tissues** | **31 organs** |  | **3,119 MS runs** | **498 Samples** |  |  |  |  |  |

The sample and experimental metadata was manually curated from their respective publications or by contacting the original authors/submitters. Metadata was annotated using Annotare [27] and stored using the Investigation Description Format (IDF) and Sample and Data Relationship Format (SDRF) file formats, required for their integration in Expression Atlas. The IDF includes an overview of the experimental design including the experimental factors, protocols, publication information and contact information. The SDRF file includes sample metadata and describes the relationship between various sample characteristics and the data files included in the dataset.

In addition to the quantification of proteins in healthy tissues representing baseline conditions described in this study, we also analysed samples in the same datasets that were from non-healthy/non-normal samples which were included in the same datasets (which are not discussed in this manuscript, but the results are also available in Expression Atlas). The selected datasets are listed in Table 1, including the original dataset identifiers, tissues and organs included, number of MS runs and number of samples. The 24 datasets sum up a total of 498 samples from 67 different tissues classified in 31 organs.

### Proteomics raw data processing

Datasets were analysed separately, using the same software and search database. Peptide/protein identification and protein quantification were performed using MaxQuant [28, 29] (version 1.6.3.4), on a high-performance Linux computing cluster. The input parameters for each dataset such as MS1 and MS2 tolerances, digestive enzymes, fixed and variable modifications were set as described in their respective publications together with two missed cleavage sites. PSM (Peptide Spectrum Match) and protein FDR (False Discovery Rate) levels were set at 1%. Other MaxQuant parameter settings were left as default: maximum number of modifications per peptide: 5, minimum peptide length: 7, maximum peptide mass: 4,600 Da. For match between runs, the minimum match time window was set to 0.7 seconds and the minimum retention time alignment window was set to 20 seconds. The MaxQuant parameter files are available for download from Expression Atlas. The UniProt human reference proteome release-2019_05 (including isoforms, 95,915 sequences) was used as the target sequence database. The inbuilt MaxQuant contaminant database was used and the decoy database were generated by MaxQuant at the time of the analysis (on-the-fly) by reversing the input database sequences after the respective enzymatic cleavage. The datasets were run in a multithreading mode with a maximum of 60 threads and 300 GB of RAM per dataset.

### Post-processing

The results coming from MaxQuant for each dataset were further processed downstream to remove potential contaminants, decoys and protein groups which had fewer than 2 PSMs. The protein intensities were normalised using the Fraction of Total (FOT) method, wherein each protein “iBAQ” intensity value is scaled to the total amount of signal in a given MS run and transformed to parts per billion (ppb).

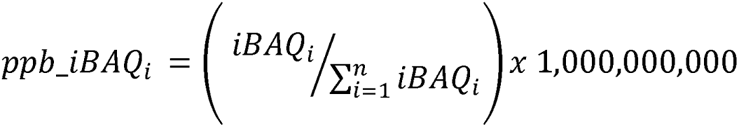

The bioconductor package ‘mygene’ [30] was used to assign Ensembl gene identifiers/annotations to the protein groups by mapping the ‘majority protein identifiers’ within each protein group. This step is required for integration into Expression Atlas, because at present, all abundance values have to be in the same reference system to be integrated. The protein groups, whose protein identifiers were mapped to multiple Ensembl gene IDs, were not integrated into Expression Atlas, but are available in Supplementary Table 1. In the case of a protein group containing isoforms from the same gene, these mapped to a single unique Ensembl gene ID and were not filtered out. In cases where two or more protein groups mapped to the same Ensembl gene ID, their median intensity values were considered. The parent genes to which the different protein groups were mapped to are equivalent to ‘canonical proteins’ in UniProt (https://www.uniprot.org/help/canonical_and_isoforms) and therefore the term protein abundance is used to describe the protein abundance of the canonical protein throughout the manuscript.

### Integration into Expression Atlas

The calculated canonical protein abundances (mapped as genes), the validated SDRF files and summary files detailing the quality of post-processing were integrated into Expression Atlas (release 37, March 2021) as proteomics baseline experiments (E-PROT identifiers are available in Table 1).

### Protein abundance comparison across datasets

Since datasets were analysed separately, the protein abundances, available in ppb values within each dataset were converted into ranked bins for comparison of abundances across datasets. The normalised protein abundances per MS run, as described above, were ranked and grouped into 5 bins, wherein proteins with the lowest protein abundance values were in bin 1 and those with the highest abundance values were in bin 5. Additionally, distinct tissue regions or organs within a dataset were grouped into batches and were binned separately. In this study, ‘tissue’ is defined as a distinct functional or structural region within an ‘organ’. For example, corpus callosum, anterior temporal lobe, dorsolateral prefrontal cortex were defined as tissues that are part of the brain (organ) and similarly left ventricle, aorta and tricuspid valve are defined as tissues in heart (organ).

During the rank-bin transformation, if a protein was not detected in any of the samples within a batch, we did not assign it a bin value, but annotated it as an NA (corresponding to not detected) value instead. However, if a protein was not detected in some samples of the batch but had protein abundance values in other samples within the batch, we assigned the lowest bin value 1 to those samples in that batch that were undetected. For example, in a dataset comprising tissue samples from brain, all samples from tissue regions such as corpus callosum, were grouped into a batch and the ppb abundances were transformed into bins. If any of the samples within a batch had no abundance values for a protein, they were marked as NA. If some samples within the batch had missing abundance values, the missing abundance values of those samples for that protein were assigned the bin value 1. Binned abundances of those proteins that were detected in at least 50% of the samples in heart and brain datasets were selected for PCA (Principal Component Analysis). To compare which normalisation methods performed better at removing batch effects, the iBAQ protein abundances were also normalised using the ComBat [31] and Limma [32] methods. PCA was performed in R using the Stats package. A Pearson correlation coefficient for all samples was calculated on pairwise complete observations of bin transformed iBAQ values in R. Samples were hierarchically clustered on columns and rows using Euclidean distances.

### Comparison of the results with the protein abundance values from the Human Protein Atlas and ProteomicsDB

Results from our analysis were compared with protein abundance data available at the HPA. Abundance profiles of proteins in normal human tissues were downloaded from HPA version 21.0. Protein abundance with reliability score labelled as ‘Uncertain’ were not considered in the comparison. For the purposes of easing the comparison and computing correlation, the categorical protein abundance levels in data downloaded from the HPA were assigned numerical values closely matching the protein abundance bins used in our analysis. Protein abundance levels annotated as ‘Low’, ‘Medium’ and ‘High’ were assigned values 1, 2 and 3 respectively. The level annotated as ‘Not detected’ was assigned NA and those annotated as ‘Ascending’, ‘Descending’ and ‘Not representative’ were all assigned a value of 1. For the purpose of this comparison, we re-binned our protein abundance data into just 3 categories: bins 1, 2, and 3 representing low, medium and high abundances, respectively. The ‘randomised edit distance difference’ was calculated across all pairs of organs included in this study and HPA. The ‘randomised edit distance difference’ is the difference between the ‘true edit distance’ and the ‘randomised edit distance’ of protein abundance bins. Randomised edit distance difference = mean(random edit distance_1-n_ - true edit distance_1-n_). The ‘true edit distance’ of a protein was computed as the absolute difference between the protein abundance bins of both pairs. The ‘randomised edit distance’ is calculated as the mean of the absolute difference between bin value of pair 1 and randomised bin value of pair 2, after sampling it 10 times, ie., randomised edit distance = mean 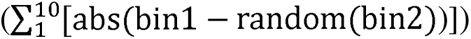. This was done using the base R package.

Normalised protein intensities from ProteomicsDB [33] were queried for organs that were in common in our study (31 organs). Values were obtained using ProteomicsDB Application Programming Interface. For different tissue samples we aggregated the normalised intensities using the median of their respective organs. The intensities were log2 normalised and compared.

### Comparison of label-free protein abundances with protein abundances generated using a TMT approach

The protein abundances calculated across various baseline human organs/tissues using the TMT-labelling method were obtained from [3] (Supplementary file ‘NIHMS1624446-supplement-2’, sheet: ‘C protein normalized abundance’). Protein abundances of the respective organs measured across different TMT channels and runs were aggregated using the median and log2 transformed. Different tissue samples from oesophagus, heart, brain and colon were aggregated into their respective organs. Pearson’s correlation was calculated in R.

### Organ-specific expression profile analysis

To investigate the organ-specific protein-based abundance profile, we carried out a modification of the classification scheme done by Uhlén *et al.* [4]. Briefly, each of the 13,070 canonical proteins that were mapped from the protein groups, was classified into one of three categories based on the bin levels in 31 organs: (1) “Organ-enriched”: one unique organ with bin values 2-fold higher than the mean bin value across all organs; (2) “Group enriched”: group of 2-7 organs with bin values 2-fold higher than the mean bin value across all organs; and (3) “Mixed”: the remaining canonical proteins that are not part of the above two categories.

Enriched gene ontology (GO) terms analysis was performed by means of the over-representation test, combining the “Organ-enriched” and “Group enriched” mapped gene lists for each organ. The computational analysis was carried out in the R environment with the package clusterProfiler [34] version 3.16.1, using the function enrichGO() for the GO over-representation test, using the parent gene list of all detected canonical proteins as the background set. Setting the p-value cut-off to 0.05 and the q-value cut-off to 0.05. Additionally, Reactome [35] pathway analysis was carried out by using mapped gene lists (indicated by the protein groups) and running pathway-topology and over-representation analysis. First, “Project to human” option was selected with the combining list of “Organ-enriched” and “Group enriched” entities. Afterwards, those pathways with p-value > 0.05 were filtered out. The hierarchical clustering was done based on the distances calculated on the p-values using the ggdendro package in R.

## Results

### Human baseline proteomics datasets

We manually selected 24 label-free publicly available human proteomics datasets coming from PRIDE and from the AMP-AD Knowledge Portal databases (Table 1). These datasets were selected to represent baseline conditions and therefore included samples annotated as healthy or normal from a wide range of biological tissues. The datasets were restricted to include those label-free datasets generated on Thermo Fisher Scientific Instruments. See more details about dataset selection in the ‘Methods’ section.

In total the aggregated datasets represent 67 healthy tissues, corresponding to 3,119 MS runs covering 498 samples, coming from 489 individuals. In this study, ‘tissue’ is defined as a distinct functional or structural region within an ‘organ’. The cumulative CPU time used for the reanalyses was approximately 2,750 hours or 114 calendar days. The numbers of protein groups, peptides, unique peptides identified and protein coverage in each dataset are shown in Table 1.

The resulting protein abundances of all samples have been made available in Expression Atlas. These ‘proteomics baseline’ quantification results can be viewed as abundance heatmaps against the gene symbols and the quantification matrices can be downloaded as text files together with annotated metadata of donor samples, experimental parameters, and a summary file describing the analysis with representative charts (quality assessment) summarising the output of the post-processed samples. The protocol for data reanalysis is summarised in Figure 1.

**Figure 1.**
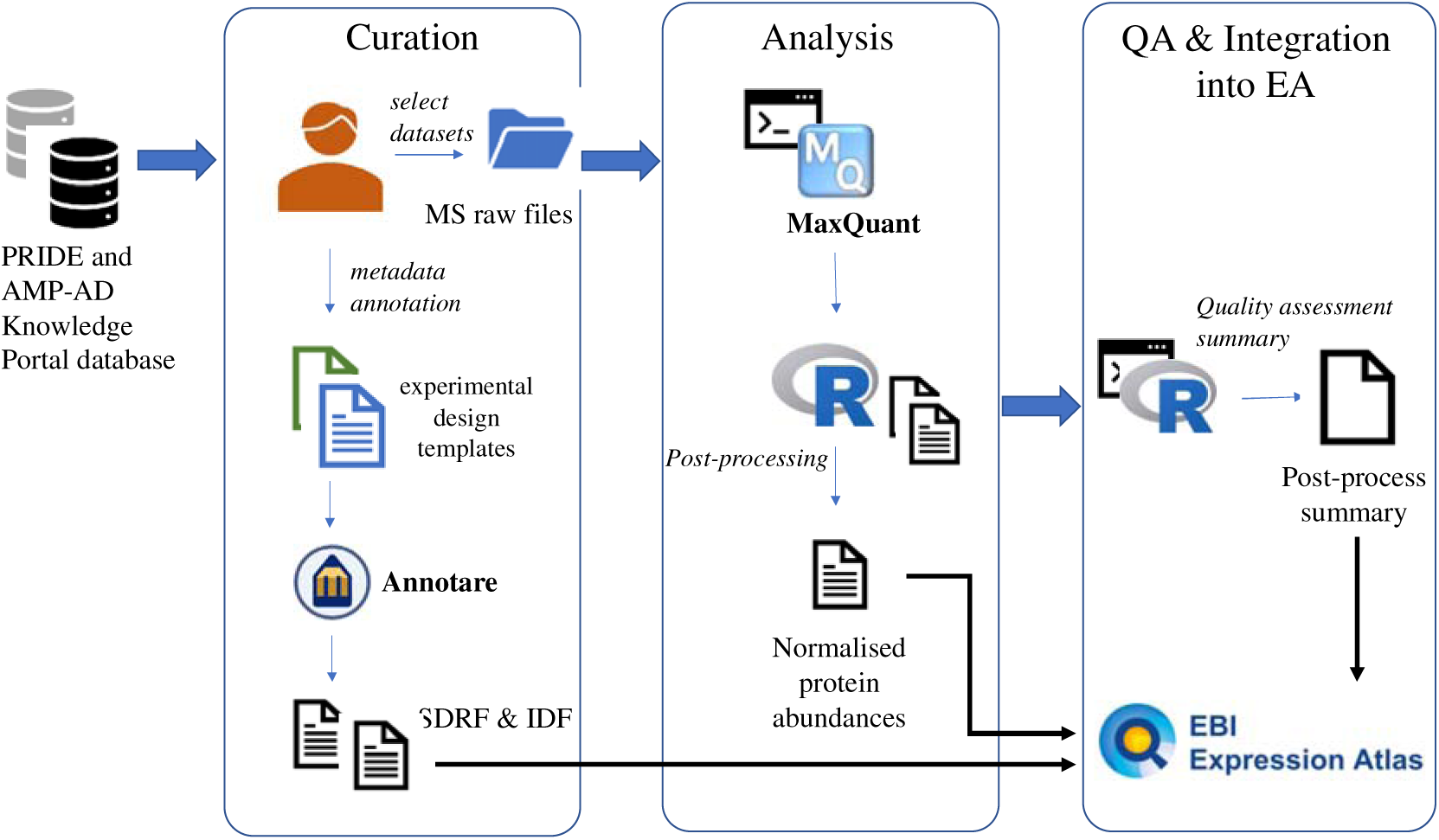
An overview of the study design and reanalysis pipeline. QA: Quality assessment.

### Protein coverage across samples

For simplicity of comparison, we broadly grouped 67 tissues into 31 major types of organs. As explained in ‘Methods’, we defined ‘tissue’ as a distinct functional or structural region within an ‘organ’. For example, corpus callosum, anterior temporal lobe, dorsolateral prefrontal cortex were all defined as tissues in brain (which is the ‘organ’). After post-processing the output files from MaxQuant, 11,653 protein groups (36.3% of identified protein groups across all datasets) were uniquely present in only one organ and 380 protein groups (1.2%) were ubiquitously observed (Supplementary Table 2). This does not imply that these proteins are unique to these organs. Merely, this is the outcome considering the selected datasets.

We mapped the isoforms in the protein groups to their respective parent gene names, which we will use as equivalent to ‘canonical proteins’ in UniProt (see ‘Methods’), from now on in the manuscript. Overall, 13,070 different genes were mapped from protein identifiers in the protein groups. We denote the term ‘protein abundance’ to mean ‘canonical protein abundance’ from here on. We then estimated the number of proteins identified across organs, which indicated that greater than 70% of all canonical proteins were present in a majority of organs (Figure 2A, 2C). We also observed the highest numbers of common proteins in samples from tonsil (92.2%) and brain (90.9%) and the lowest numbers in samples from umbilical artery (7.2%).

**Figure 2.**
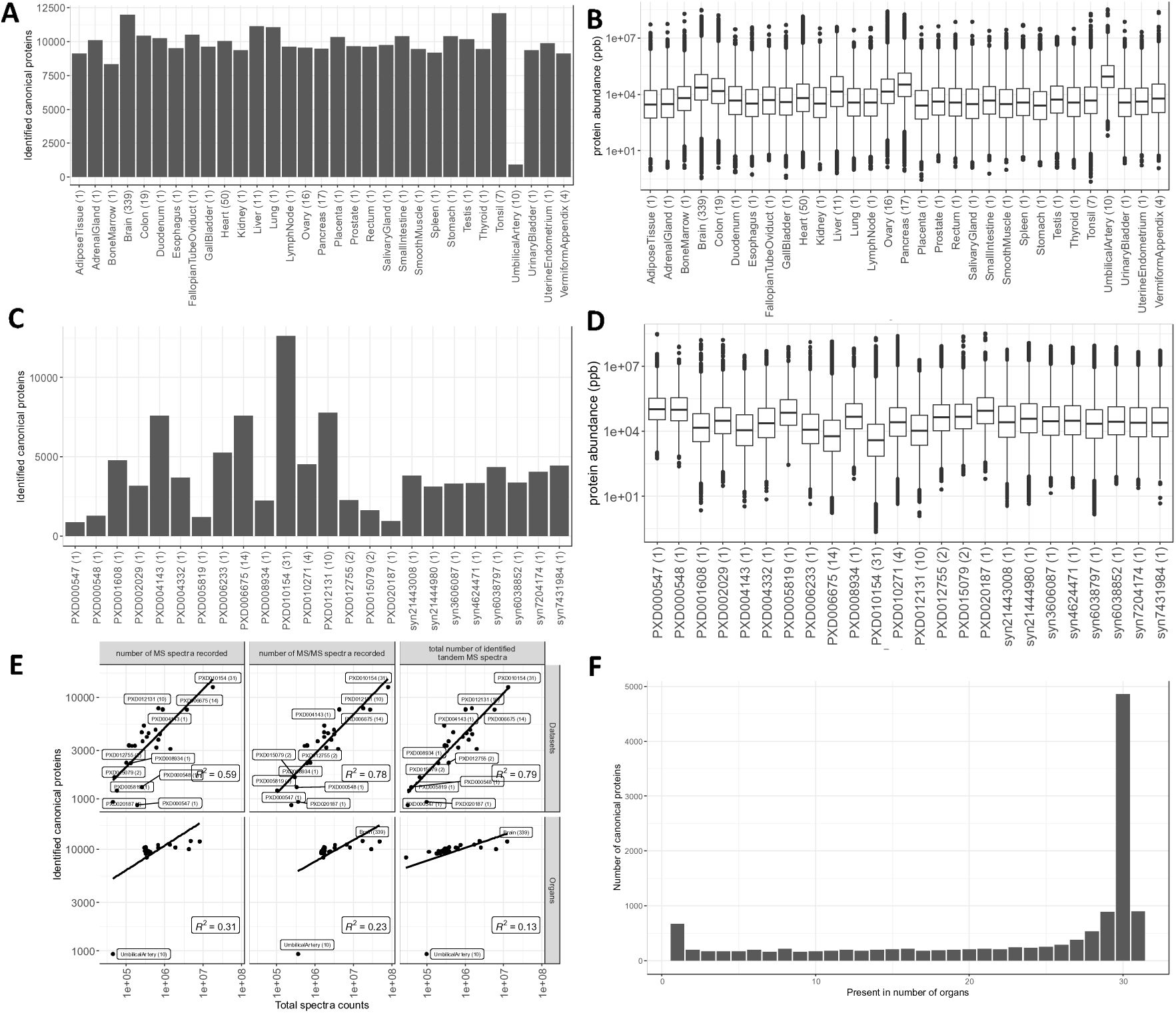
(A) Number of canonical proteins identified across different organs. The number within the parenthesis indicates the number of samples. (B) Range of normalised iBAQ protein abundances across different organs. The number within the parenthesis indicates the number of samples. In Panels (A) and (B), the term heart is used in a broader sense to mean cardiovascular system. (C) Canonical proteins identified across different datasets. The number within the parenthesis indicate the number of unique tissues in the dataset. (D) Range of normalised iBAQ protein abundances across different datasets. The number within parenthesis indicate the number of unique tissues in the dataset. (E) Comparison of total spectral data with the number of canonical proteins identified in each dataset and organ. (F) Distribution of canonical proteins identified across organs.

The higher number of proteins identified in brain could be attributed to the greater representation of samples (339 samples out of 498, 68.0%). However, tonsil was represented only by 7 samples and were all derived from one dataset (PXD010154). It is worth noting that the sample preparation protocol for the tonsil samples employed seven different proteases (Trypsin, LysC, ArgC, GluC, AspN, LysN and Chymotrypsin) for tissue digestion [36], thus significantly increasing its peptide coverage [36]. The sample size of umbilical artery, which showed significantly lower protein coverage than other organs, were 10 samples.

The largest number of canonical proteins were identified in dataset PXD010154 (Figure 2C), which comprises numerous tissue samples (31 tissues) including samples from tonsil. The dynamic range of protein abundances in all organs is shown in Figure 2B. On the other hand, protein abundances among datasets showed that PXD010154 had the lowest median protein abundances (Figure 2D). We also compared the quantity of spectral data from various organs and datasets with the number of canonical proteins identified in them, to detect any organ or dataset that showed enrichment of proteins relative to the amount of data. We observed a linear relation between the number of proteins identified and the amount of spectral data present in the organ samples or datasets (Figure 2E).

### Distribution of canonical protein identifications per organ

We observed that 37.1% (4,853) of the identified canonical proteins were expressed in 30 different organs (Figure 2F). The low number of proteins identified in umbilical artery (933) samples greatly influenced the protein distribution. As a result, 7.0% (917) of all identified canonical proteins were present in all 31 organs, whereas 4.2% (565) of the identified canonical proteins were present uniquely in one organ. However, it is important to highlight that the list of concrete canonical proteins that were detected in just one organ should be taken with caution since the list is subjected to inflated FDR, due to the accumulation of false positives when analysing the datasets separately. However, this should not be an issue in the case of proteins detected across 5 datasets or more, since the number of commonly detected decoy protein hits enabled to calculate a protein FDR less than 1% (Figure S1 in Supplementary Figures).

### Protein abundance comparison across organs

Next, we compared the protein abundances to see how proteins compared across different organs. Inter-dataset batch effects make comparisons challenging. We transformed the normalised iBAQ intensities into ranked bins as explained in ‘Methods’. The bin transformed protein abundances in all organs are provided in Supplementary Table 3.

To compare protein abundance across all organs, a pairwise Pearson correlation coefficients of binned protein abundances was calculated across 498 samples (Figure 3). We observed a good correlation of protein abundance within the brain (median R^2^ = 0.61) and cardiovascular system (median R^2^ = 0.41) samples, which represent the two organ groups with the largest number of samples. We tested the effectiveness of various normalisation methods in reducing batch effects, by performing a PCA on samples coming from cardiovascular system and brain datasets. The brain and cardiovascular system samples analysed constituted the largest numbers in the aggregated dataset, including 19 and 3 datasets, respectively. First, we performed PCA on the normalised iBAQ values, wherein the brain samples did not cluster either by tissues or by datasets. However, for cardiovascular system samples, we observed clustering of samples by datasets and not by tissue type (Figure S2 in Supplementary Figures). We then tested the ComBat and Limma normalisation methods on iBAQ values, which neither showed clustering of samples by tissues nor by datasets for both cardiovascular system and brain samples (Figures S3 and S4 in Supplementary Figures).

**Figure 3.**
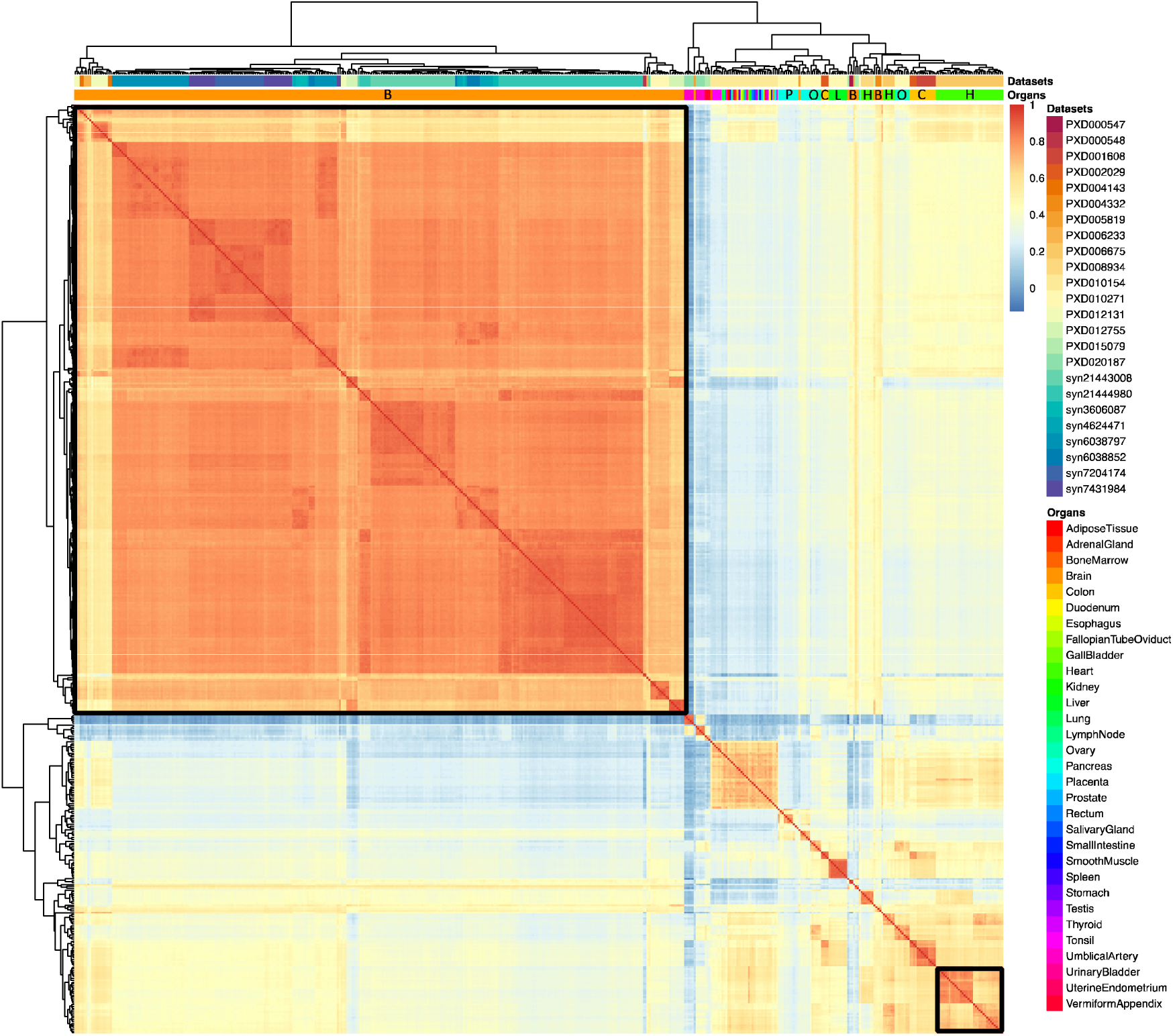
Heatmap of pairwise Pearson correlation coefficients across all samples. Colours on the heatmap represents the correlation coefficient and was calculated using the bin transformed iBAQ values. The samples are hierarchically clustered on columns and rows using Euclidean distances. The clusters composed of brain and cardiovascular system (heart) samples are highlighted with black borders. The abbreviations used in the organs’ header are: B: Brain, C: Colon, H: Heart, L: Liver, O: Ovary and P: Pancreas.

We then decided to use the bin-transformed protein abundances (see ‘Methods’). First, we observed that brain samples were clustered together according to their tissue type (Figure 4A). All brain tissue samples, except those coming from the dorsolateral prefrontal cortex (DLPFC) were part of individual datasets. The DLPFC samples were derived from six separate datasets, of which five of them were part of the Consensus Brain Protein Coexpression study [26]. The DLPFC samples clustered into two groups: a large group that comprised samples from the Consensus Protein Coexpression study and a smaller cluster with samples from dataset PXD004143 (Figure 4B), indicating that there was still a residual batch effect.

**Figure 4.**
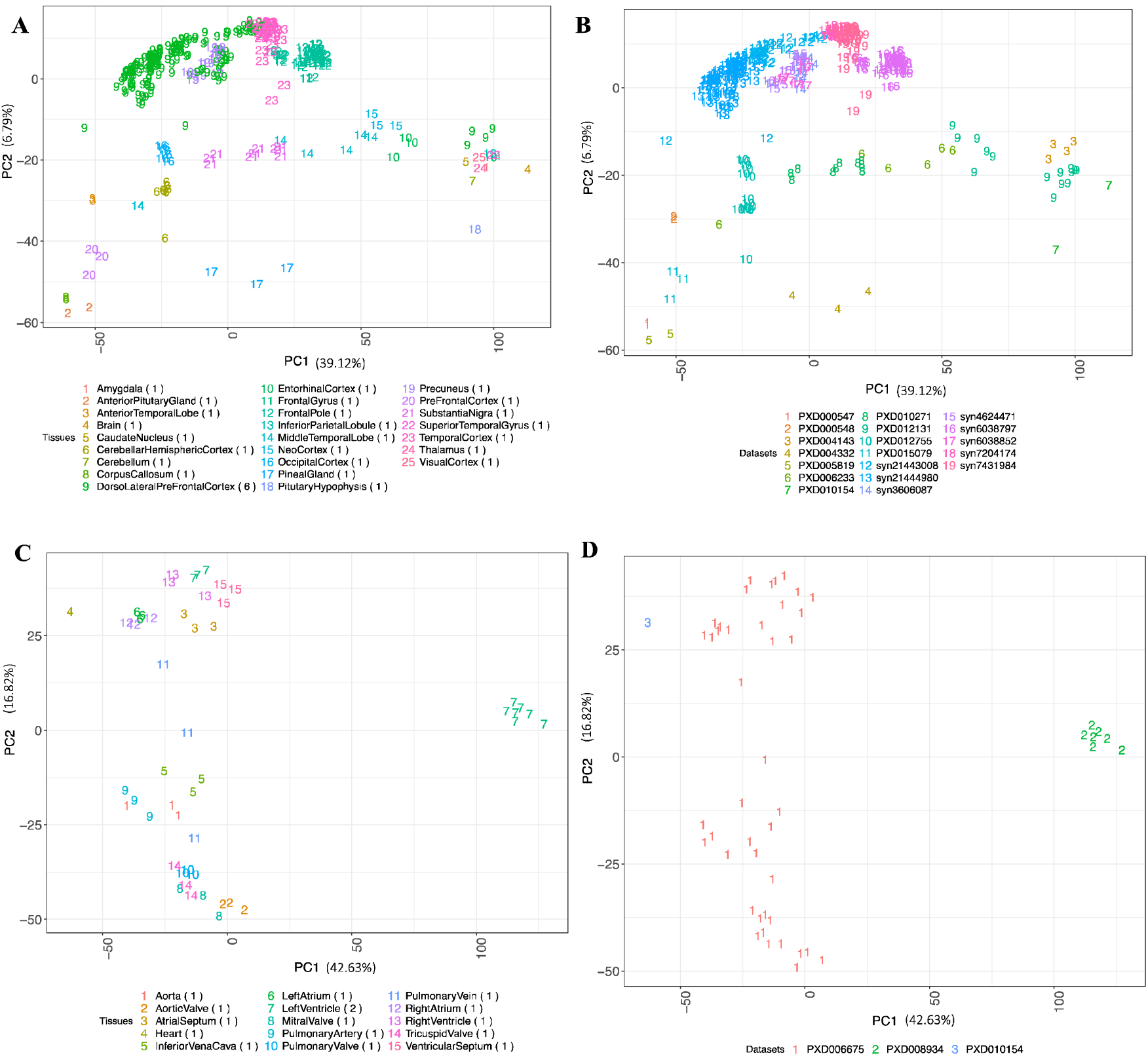
(A) PCA of brain samples coloured by the tissue types. (B) PCA of brain samples coloured by their respective dataset identifiers. (C) PCA of cardiovascular system (heart) samples coloured by the tissue types. (D) PCA of cardiovascular system (heart) samples coloured by their respective dataset identifiers. The numbers in parenthesis indicate the number of datasets for each tissue. Binned values of canonical proteins quantified in at least 50% of the samples were used to perform the PCA.

Similarly, we observed cardiovascular system samples clustered according to their tissue types (Figure 4C). All cardiovascular system samples except those coming from left ventricle were part of an individual dataset. Interestingly, we observed 3 major clusters: one wherein all valve samples (aortic valve, mitral valve, pulmonary valve and tricuspid valve) were clustered together. A second cluster where the samples from ventricles and atriums were clustered in a large group together with other cardiovascular system samples. Finally, left ventricle samples from dataset PXD008934 (Figure 4D) formed a separate cluster indicating that there were still batch effects which were not completely removed.

### Comparison of protein abundance values with previous studies

We first compared the protein abundances resulting from our reanalysis with those reported in the original publications. By comparing the number of protein groups or genes identified in individual datasets we observed that the differences between our analysis and the original published results ranged from as low as 1.3% (E-PROT-53, dataset syn7204174) to as high as 43.2% (E-PROT-36, dataset PXD012755). Similarly, the difference at the level of identified peptides ranged from a minimum of 0.29% (E-PROT-33, dataset PXD005819) to a maximum of 57.2% (E-PROT-36, dataset PXD012755) (Supplementary Table 4). These differences in overall numbers could be due to various factors, including the used target protein sequence database and the analysis software and version used.

We then compared our results with protein abundance data available in ProteomicsDB [33] and found a good correlation in abundance across various organs. As it can be seen in Figure S5 in Supplementary Figures the highest correlation was found in salivary gland (R^2^ = 0.75) and the lowest one in ovary (R^2^ = 0.52). However, it should be noted that one of the datasets included in our analysis (dataset PXD010154) is also included in ProteomicsDB. Additionally, we also made a comparison between our protein abundance results and those found in a large study across multiple human organs using TMT-labelling method [3]. Figure S6 in Supplementary Figures shows the Pearson’s correlations of protein abundances between both studies, which was generally lower than in the case of ProteomicsDB data, ranging from 0.22 to 0.48 across various organs.

In addition, we compared our results with protein abundances computed using antibody-based methods, available in the Human Protein Atlas (HPA). Firstly, we performed a qualitative analysis in which we compared the number of proteins identified in matching organs in our analysis with those proteins identified in the HPA. There were 30 organs that were in common between both studies (except for umbilical artery which was not available in HPA). The comparison results are shown in Figure S7 in Supplementary Figures. Our analysis shows that an average of 43.7% of all proteins identified in HPA were also present in our aggregated dataset, with the highest number of common identified proteins found in brain (50.4%) and the lowest number of proteins in common was in adipose tissue (27.2%). On the other hand, an average of 40.4% of proteins were only identified in our analysis and were not present in the results analysed in HPA. The largest and the lowest number of proteins that were identified only in our analysis were in adipose tissue (61.6%) and in testis (30.2%), respectively. Lastly, an average of 15.8% of the proteins were exclusive to HPA and not identified across any organs in our analysis. Of these proteins the largest HPA exclusive group of proteins was present in vermiform appendix (21.6%) and the lowest was found in adrenal gland (8.9%).

We then compared protein abundances by first transforming the abundances in HPA into numerical bins. Protein abundance data from HPA are annotated in 3 categorical groups as ‘Low’, ‘Medium’ and ‘High’, which, we converted into 3 numerical bins 1, 2, and 3 respectively. For the purpose of this comparison, we re-binned our protein abundance data into just 3 categories: bins 1, 2, and 3 representing low, medium and high abundances, respectively (see ‘Methods’). To identify difference between noise and signal we calculated the randomised edit distance difference metric across all organs between the two studies (see Methods). The higher ‘randomised edit distance difference’ indicates that there is a difference between signal and random noise. The randomised edit distance difference matrix (Figure S8 in Supplementary Figures) shows that the randomised edit distance difference between organs within HPA are low (average randomised edit distance difference = 0.18) compared to that of organs within our study (average randomised edit distance difference = 0.43). This seems to suggest that the overall protein abundances generated in this study are less noisy than the abundance data available in HPA.

### The organ elevated proteome and the over-representative biological processes

As explained in ‘Methods’, according to their abundances, canonical proteins were divided in three different groups according to their organ-specificity: “Organ-enriched”, “Group enriched” and “Mixed” (see Supplementary Table 5). We considered elevated canonical proteins those which were classified as an “Organ-enriched” or “Group enriched” instead of the “Mixed” group. The analysis (Figure 5A) showed that on average, 3.8% of the total elevated canonical proteins were organ group-specific. The highest ratio was found in the adrenal gland (9.3%), brain (7.5%) and liver (7.1%), and the lowest ratio in gall bladder (2.3%) and umbilical artery (0.1%). In addition, 0.4% of the total canonical proteins were unique organ-enriched. The highest ratio was found in brain (3.8%), cardiovascular system (1.4%) and liver (0.5%) and the lowest ratio (∼0.1%) was found in tonsil and uterine endometrium.

**Figure 5.**
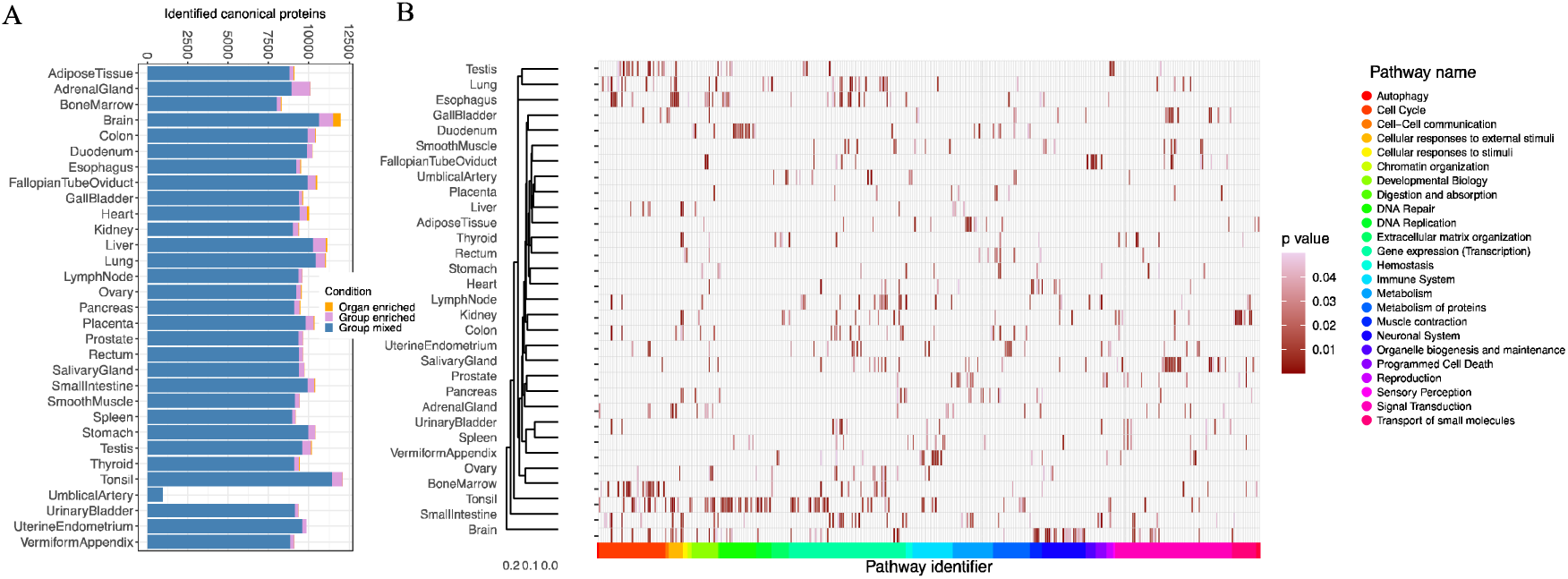
(A) Analysis of organ-specific canonical proteins. The analysis comprises the number of canonical proteins found in 31 organs, classified in three groups: “organ-enriched”, “group enriched” and “group mixed”. (B) Pathway analysis of the over-represented canonical proteins, showing the statistically significant representative pathways (p-value < 0.05) in 31 organs. In Panels (A) and (B), the term heart is used in a broader sense to mean cardiovascular system.

Then, we performed a Gene Ontology (GO) enrichment analysis using the GO terms related to biological processes for those canonical proteins that were “organ-enriched” and “group-enriched”, is shown in Table 2. As a summary, 358 GO terms were found statistically significative across all organs (see Supplementary Table 6). The terms found were in agreement with the known functions of the respective organs. The brain had the largest number of “organ-enriched” canonical proteins (457), among the biological processes associated stand out the regulatory function on membrane potential (GO:0042391), neurotransmitter transport (GO:0006836), modulation of chemical synaptic transmission (GO:0050804), regulation of trans-synaptic signalling (GO:0099177) and potassium ion transport (GO:0006813). The second organ with a greater number of “organ-enriched” canonical proteins was cardiovascular system (137). The enriched biological processes involved were related with striated muscle cell differentiation (GO:0051146), sarcomere organisation (GO:0045214), muscle structure development (GO:0061061) and regulation of myotube differentiation (GO:0010830). As expected, there were common GO terms that were shared between the organs, such as: detoxification of inorganic compound (GO:0061687) in liver and kidney, import across plasma membrane (GO:0098739) in kidney, brain and umbilical artery, processes involved in tissues with high cell division turnover like chromosome segregation (GO:0007059) in bone marrow and testis.

**Table 2.** Analysis of the GO terms for each organ using the elevated organ-specific canonical proteins and group-specific as described in the ‘Methods’ section.

| Organ | GO ID | Description | Adjusted p-value |
| --- | --- | --- | --- |
| **Adrenal gland** | GO:0031649 | Heat generation | $7.94 \times 10^{-4}$ |
| **Bone marrow** | GO:0034080 | CENP-A containing nucleosome assembly | $1.49 \times 10^{-3}$ |
| **Brain** | GO:0042391 | Regulation of membrane potential | $1.71 \times 10^{-11}$ |
| **Fallopian tube oviduct** | GO:0044782 | Cilium organization | $2.88 \times 10^{-48}$ |
| **Gallbladder** | GO:0017158 | Regulation of calcium ion-dependent exocytosis | $1.17 \times 10^{-2}$ |
| **Cardiovascular system (Heart)** | GO:0051146 | Striated muscle cell differentiation | $9.58 \times 10^{-6}$ |
| **Kidney** | GO:0046942 | Carboxylic acid transport | $4.81 \times 10^{-16}$ |
| **Liver** | GO:0097501 | Stress response to metal ion | $1.08 \times 10^{-3}$ |
| **Lung** | GO:0003002 | Regionalization | $8.83 \times 10^{-3}$ |
| **Lymph node** | GO:0002250 | Adaptive immune response | $1.93 \times 10^{-4}$ |
| **Ovary** | GO:0008544 | Epidermis development | $8.42 \times 10^{-7}$ |
| **Placenta** | GO:0044706 | Multi-multicellular organism process | $4.84 \times 10^{-3}$ |
| **Testis** | GO:0048232 | Male gamete generation | $5.00 \times 10^{-24}$ |
| **Thyroid** | GO:0098742 | Cell-cell adhesion via plasma-membrane adhesion molecules | $1.34 \times 10^{-2}$ |
| **Tonsil** | GO:0031424 | Keratinization | $3.20 \times 10^{-5}$ |
| **Umbilical artery** | GO:0001937 | Negative regulation of endothelial cell proliferation | $1.75 \times 10^{-3}$ |

Next, we performed a pathway-enrichment analysis using Reactome [35] to analyse canonical proteins that were “organ-enriched” and “group-enriched” (see Supplementary Table 7). The heatmap (Figure 5B) shows the statistically significant pathways, (p-value < 0.05) across the organs. The total number of pathways found in all the organs were 928, and the largest number of pathways was found in the brain with 67 pathways. The pathways found were consistent with the GO analysis and with the expected function in each organ. We observed a ‘cell cycle’ cluster of over-represented pathways related to bone marrow and testis (R-HSA-1640170, R-HSA-69620, R-HSA-73886, R-HSA-2500257 and R-HSA-69618), expected in high cell turnover tissues, the digestion pathway (R-HSA-192456) in pancreas and stomach, a neuronal system cluster of pathways (R-HSA-112316) in the brain, and pathways related to the transport of small molecules (R-HSA-382551, R-HSA-425407, R-HSA-425393 and R-HSA-425366) in kidney.

### Integration of results into Expression Atlas

Protein abundance results from label-free experiments across various tissues were integrated into Expression Atlas. The abundances of each protein are represented in terms of their canonical gene symbols since Expression Atlas is designed as a gene-centric resource. Proteomics results can be accessed using the link www.ebi.ac.uk/gxa/experiments/E-PROT-xx/Downloads (replacing xx with the corresponding identifier for each dataset). For each dataset, the raw unprocessed MaxQuant output files (proteinGroups.txt) are made available to download together with the input experimental parameters (mqpar.xml) to MaxQuant, as well as the metadata annotation file of each sample. We also provide a summary of the quality assessment of the results. Supplementary File 8 provides a brief manual on how to access proteomics data in Expression Atlas.

## Discussion

We here include a combined analysis of human baseline proteomics datasets representing baseline protein abundance across 67 healthy tissues grouped in 31 organs. This type of study has been enabled by the large amount of data in the public domain, as the proteomics community is now embracing open data policies. The large-scale availability of MS data in public databases such as PRIDE enables integrated metaanalyses of proteomics data covering a wide array of tissues and biological conditions. The main aim of our study was to provide a systems-wide baseline protein abundance catalogue across various tissues and organs, which could be used as a reference (especially to those non-experts in proteomics) and help to reduce redundant efforts of similar computationally expensive reanalyses.

Unlike what was done in one previous study performed by us [22], and analogously to what we did with a more recent study performed using data generated from baseline rat and mouse tissues[23], here we analysed each dataset separately using the same software and the same search protein sequence database. The disadvantage of this approach is that the FDR statistical thresholds are applied at a dataset level and not to all datasets together as a whole, with the potential accumulation of false positives across datasets. However, this does not represent an issue in the case of proteins detected in common several datasets (in this particular study, at least 5 datasets will provide a protein FDR of less than 1%, Figure S1 in Supplementary Figures), since the number of commonly detected false positives is reduced in parallel with the increase in the number of common datasets where a given protein is detected. This means that proteins that are only detected in a small number of datasets could potentially be false positives (considering the applied 1% FDR at the protein level), but that does not mean that they are. At that point, researchers should seek for confirmation of the existence of the protein (if that is their goal) via alternative sources as well. Different reanalyses of some of the datasets used in this study, with different FDR calculation methods, have been published independently [56, 57].

In our view, the objective of integrating quantitative proteomics information with other omics data types (in this case transcriptomics) in resources used by non-proteomics researchers such as Expression Atlas is only feasible in a sustainable manner using a dataset per dataset analysis approach, at least at present. This enables that: (i) computing requirements for the reanalyses are realistic given the large volume of files included in the potentially very large-combined datasets; (ii) interesting additional datasets could be added at a different time point without having to reanalyse all datasets together again; (iii) future updates in the results are more feasible to perform; and (iv) (semi)-automation of the reanalyses is achievable, making again these efforts more sustainable. We followed this same overall approach in the recent study we performed in mouse and rat tissues in baseline conditions [23]. Additionally, we compared our results with previous analogous studies performed in baseline tissue using MS and also the antibody-based data available in the HPA. These comparisons generated quite different results depending on each study.

One of the major bottlenecks was, as reported before, the curation of dataset metadata, consisting in mapping files to samples and biological conditions. Detailed sample and donor metadata is crucial for result reproducibility and we found detailed metadata available in PRIDE for just a handful of datasets. The required information either was inferred or were requested by contacting the respective study’s authors. If no responses were obtained, such datasets could not be considered for the reanalysis. Therefore, to aid reproducibility of results in the future, we need to improve the provision of metadata by data submitters. A format to enable that has been developed (the SDRF-Proteomics format, as part of the new MAGE-TAB-Proteomics format), which can be submitted optionally to PRIDE [58]. We expect that it will become increasingly used for data submissions to PRIDE, once the right tooling is available and submitters have been educated appropriately.

Another one of the major challenges in the reanalyses of a large number of proteomics datasets is the integration of results from different datasets since batch effects are inevitable. We used a rank-binned normalisation of abundances, which transformed protein abundances across datasets and samples to bins of 1 to 5. This approach is useful to reduce batch effects, although we acknowledge there is also loss of signal through this transformation. We also acknowledge that this method is not ideal in all circumstances, but in our view, it generally works better when compared to popular methods to reduce batch effects such as ComBat and Limma. Since our method computes protein abundances in terms of their canonical protein and gene identifiers, we acknowledge that using median of intensities to aggregate abundances over protein groups with isoforms coming from the same canonical protein may not represent the total dum of all proteins and may influence ranking during binning.

Although the combined dataset contains a higher representation of particular tissues (especially brain), we believe it represents the current state of the art with regard to public baseline human proteomics studies carried out in tissues. The analysis search strategy used in this study focused only on detecting known coding protein sequences, using the UniProt reference proteome, in the same way as performed in the original studies. Therefore, it was not possible to detect any Single Amino Acid Variants or equivalent isobaric combinations involving PTMs. However, the effect of this limitation in the analysis should be in our view, relatively small, because of the type of samples used in this study (healthy tissues) which did not involve e.g. tumour samples. The availability of the results through Expression Atlas enables the integration of mRNA and proteomics abundance information, offering an interface for researchers to access this type of information. The next step will be the integration of datasets in the differential part of Expression Atlas. The work required is more complex there at different levels, including the downstream statistical differential analysis. Also, availability of the mapping between the channels (e.g. in TMT, SILAC experiments) and the samples is very rare at present. In parallel, work has also started in integrating in Expression Atlas proteomics data generated using Data Independent Acquisition (DIA) approaches [59].

The generated baseline protein abundance data can be used with different purposes. For instance, quantitative proteomics data can be used for the generation of co-expression networks and/or the inference of protein complexes. Protein abundance data could also be used to potentially refine the recently developed AlphaFold-based protein complexes predictions [60]. Additionally, it is possible to use artificial intelligence approaches to impute protein abundance values using calculated abundance values as training data [61]. It would also be possible to perform expression correlation studies between gene and protein expression information. However, this type of studies can only be performed optimally if the same samples are analysed by both techniques, as reported in the original publication for dataset PXD010154 [36]. It should also be highlighted that a growing number of studies are using non-MS based proteomics techniques such as the use of affinity reagents (e.g. the Olink^®^ and Somalogic^®^ platforms), due to the increased throughput that they can provide. Initial studies are being performed to compare these with MS approaches.

In conclusion the results presented here represent a large-scale meta-analysis of public human baseline proteomics datasets. We also show the challenges in this kind of analyses, providing a roadmap for such future studies.

## Authors’ contributions

AP, DGS, SW, DJK selected and curated the datasets. AP, DGS and SW performed analyses. AC and AJ helped in the interpretation of results and designed approach for data normalisation. NG, PM and IP helped integration of results into Expression Atlas. AP, DGS, JAV wrote the manuscript. All authors have read and approved the manuscript.

## Data availability

Expression Atlas E-PROT identifiers, and PRIDE and AMP-AD original dataset identifiers are included in Table 1.

## Acknowledgements

First of all, we would like to thank all data submitters who made their datasets available in the public domain (most of the datasets in PRIDE). This work has been funded by Open Targets (project OTAR-043), Wellcome Trust [grant number 208391/Z/17/Z], BBSRC [BB/T019670/1 and BB/T019557/1] and EMBL core funding. We thank Thawfeek Varusai for helping with the pathway analysis using Reactome. We are very grateful to Mathias Walzer and Yasset Perez-Riverol for their useful suggestions and discussions.

## Abbreviations

AD: Alzheimer’s Disease
DLPFC: Dorsolateral PreFrontal Cortex
FOT: Fraction Of Total
HPA: Human Protein Atlas
GO: Gene Ontology
iBAQ: intensity-based absolute quantification
IDF: Investigation Description Format
MS: Mass Spectrometry
PCA: Principal Component Analysis
SDRF: Sample and Data Relationship Format

## Supplementary Material

**Supplementary Table 1:** Protein groups from all datasets that are mapped to more than one Ensembl Gene ID.

**Supplementary Table 2:** Median protein abundances (in ppb) for each protein group across various tissue samples in each organ.

**Supplementary Table 3:** Median binned protein abundances across various organs.

**Supplementary Table 4:** Table showing the comparison of protein and peptide identification numbers across various datasets with the reported ones in their respective original publications.

**Supplementary Table 5:** ‘Organ-enriched’ and ‘Group-enriched’ elevated proteomes in various organs.

**Supplementary Table 6:** Gene Ontology enrichment analysis of ‘organ-enriched’ and ‘group-enriched’ proteins.

**Supplementary Table 7:** Reactome pathway-enrichment analysis of “organ-enriched” and “group-enriched” proteins.

**Supplementary File 8:** Tutorial on how to browse proteomics abundance data in Expression Atlas.

**Supplementary Figures:** Supplementary figures of ComBat and Limma normalisation methods and comparison of the results with previous studies.

## Notes

### Competing Interest Statement

The authors have declared no competing interest.

### Summary of Updates

We have modified Figure 3. Heatmap of pairwise Pearson correlation coefficients across all samples, to change the colour scale of the legends and included annotations to make identification of organs easier. We have also added two additional publication references.

