## Supplemental File 8 for "An integrated view of baseline protein expression in human tissues"

### Browse proteomics experiments in Expression Atlas

You can browse through the experiments in Expression Atlas in the **Browse experiments** tab, which shows you a table listing all the experiments currently available in Expression Atlas. You can filter and/or re-order the table content using the categories and search boxes in the header line.

Proteomics experiments can be listed by typing the keyword proteomics in the **search all columns** field.

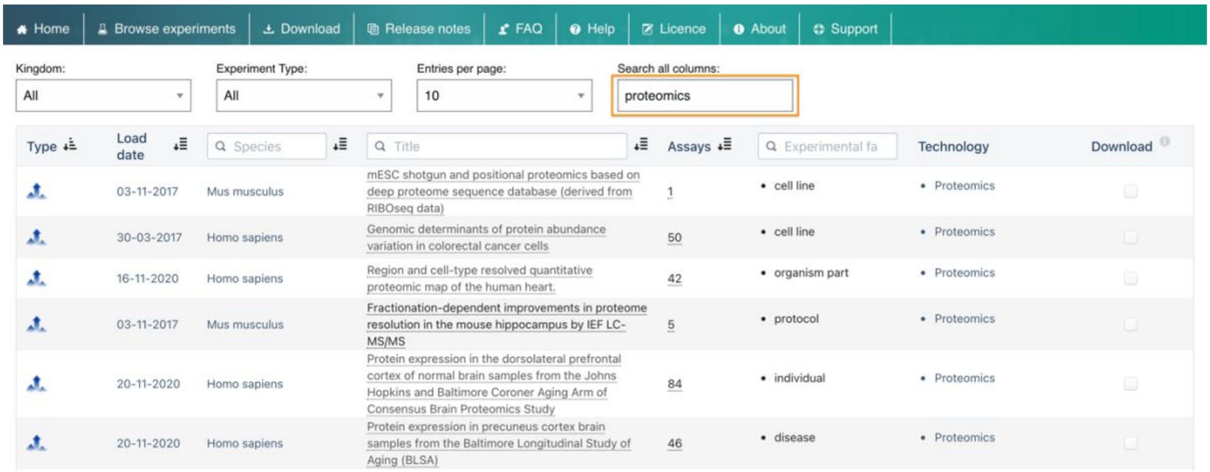


*Finding an individual dataset*

To find an individual experiment, change the experiment accession (example E-MTAB or E-PROT) in the URL [www.ebi.ac.uk/gxa/experiments/E-PROT-XX/Results](www.ebi.ac.uk/gxa/experiments/E-PROT-XX/Results). Replace E-PROT-XX with the proteomics accession of choice (for example E-PROT-52) in the URL.

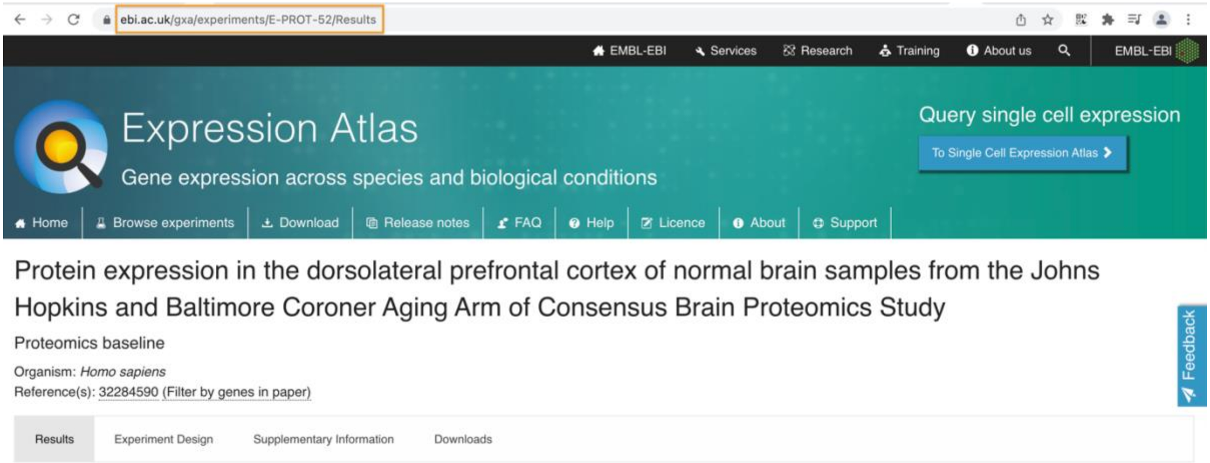

#### Baseline proteomics experiment page

Each baseline experiment in Expression Atlas has its own Experiment page.

In a baseline experiment page, expression levels are displayed in one heatmap by colour intensity, according to the gradient bar above the heatmap. The gradient shows intensities corresponding to expression levels for the 50 genes displayed. Mouse over a cell in the heatmap to see expression values for each gene in each tissue (or other condition).

For proteomics experiments the expression values are displayed as parts per billion (ppb). The default minimum expression value is 0.

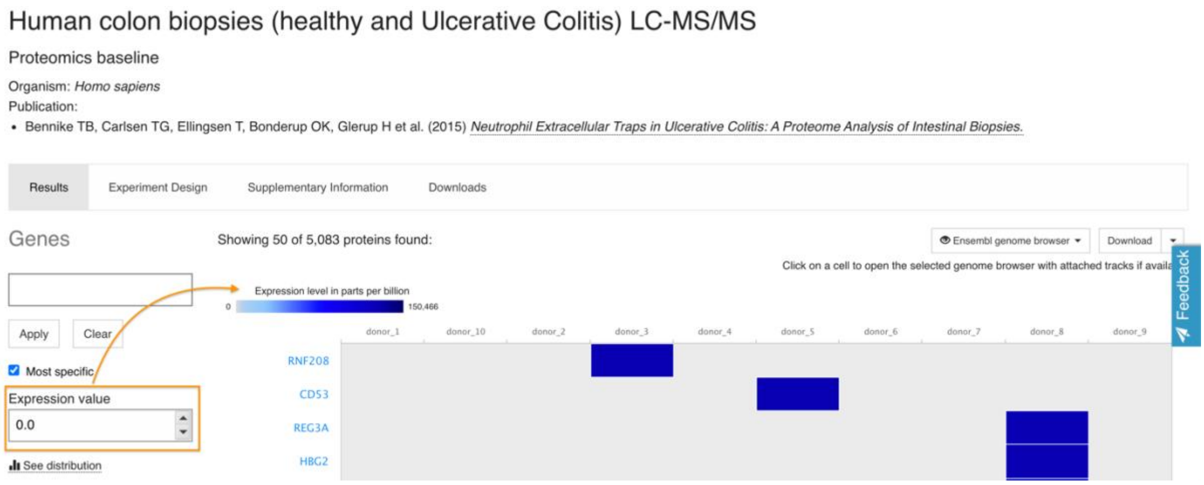


*Most specific search*

By default, the 50 most specifically expressed genes (rows) across all conditions (columns) studied are displayed. Unclick the **Most specific** option to show genes with highest expression first.

*Protein to Gene mapping and quantification*

For Mass Spectrometry baseline proteomics experiments the protein abundances are quantified in units of parts per billion (ppb). The abundances of proteins are displayed in terms of their parent gene identifiers. Expression Atlas uses a Gene ID reference frame, therefore to integrate proteomics results the UniProt protein accessions were mapped to Ensembl Gene identifiers using the bioconductor package 'mygene'. Further details are described in the 'Analysis Method' section of the Supplementary Information to each experiment.

*Other information in the baseline experiment page*

The **Experiment Design** tab shows mass spectrometry runs along with their corresponding biological sample characteristics and experimental variables values.

The **Supplementary Information** tab for Mass Spectrometry proteomics experiments the analysis methods include data processing protocols for raw Mass Spectrometry data and for post-processing the results, which also includes mapping UniProt protein accessions to Ensembl Gene identifiers.

The **Downloads** tab for Mass Spectrometry based proteomics experiments the files that one can download are: i) raw unprocessed output for baseline Data Dependent Analysis (DDA) experiments, ii) post-processed expression values, iii) quality assessment summary of the experimental runs, iv) input parameters to process raw data files for DDA experiments and v) the experimental design template of all samples.

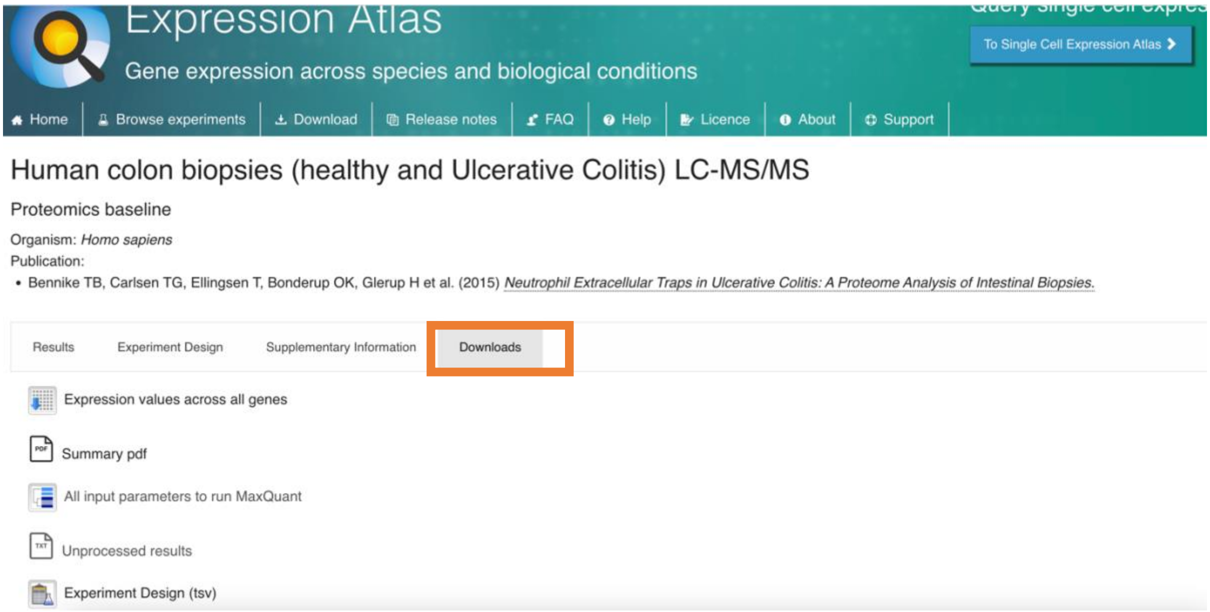
