## Supplemental Figures for "An integrated view of baseline protein expression in human tissues"

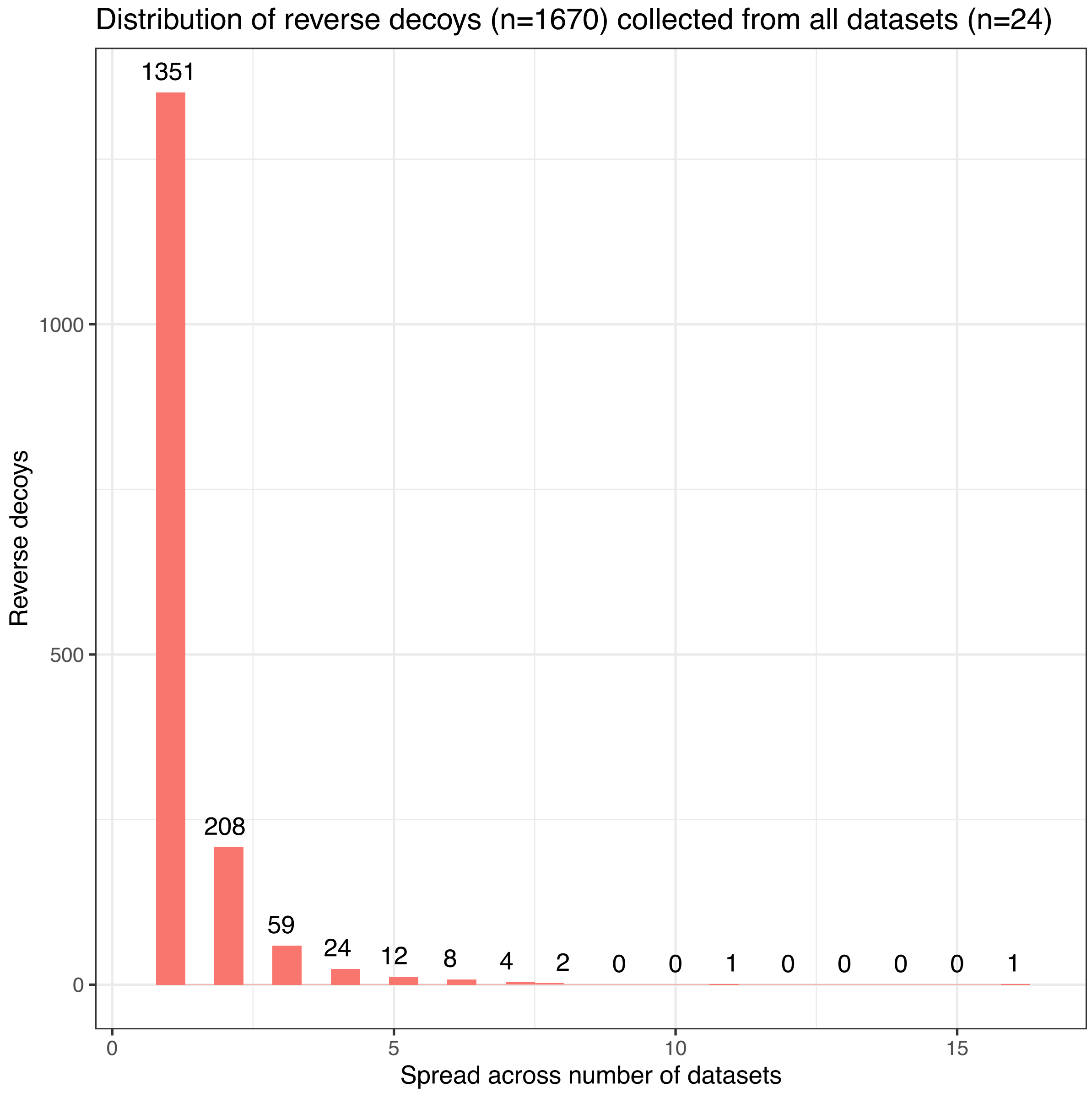


**Figure S1**. Distribution of common reverse protein decoy hits across the number of datasets.

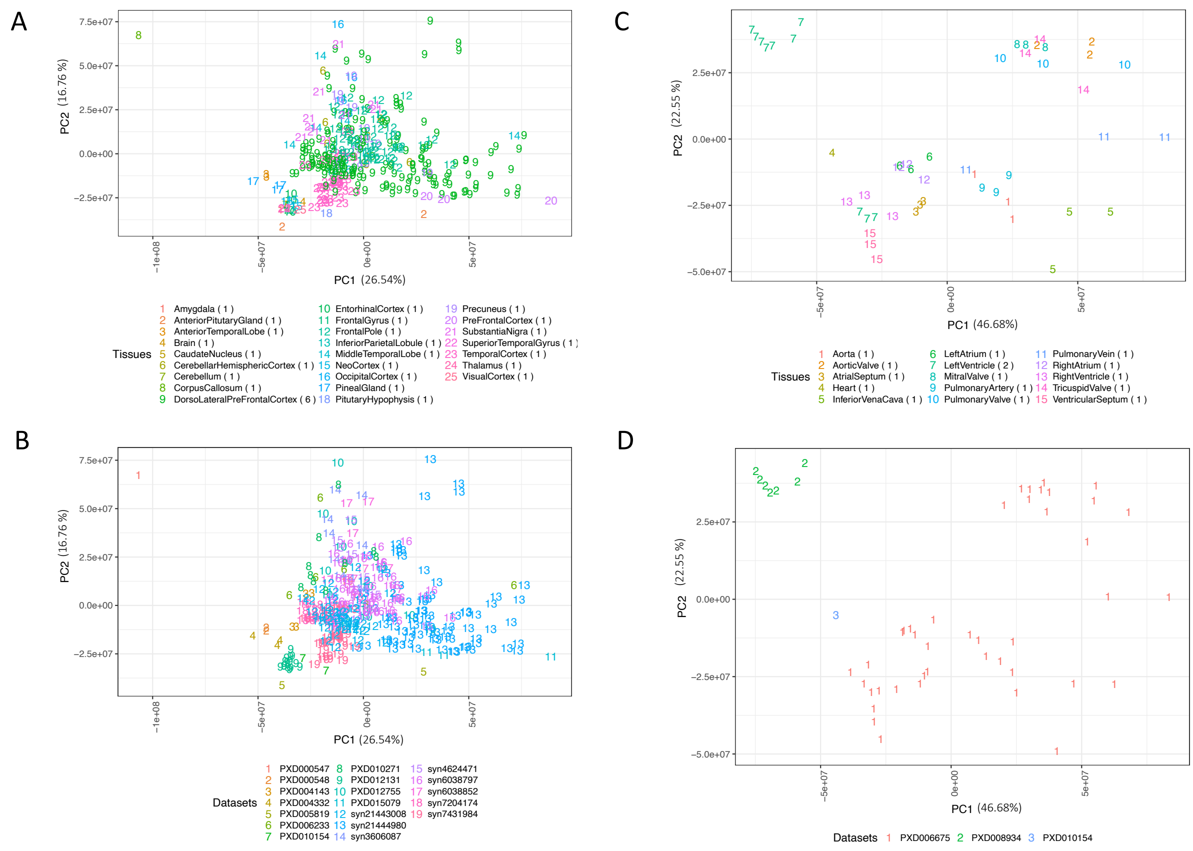


**Figure S2.** PCA of FOT normalised iBAQ protein abundances (samples without bin transformation) of (A) brain samples coloured by tissues and (B) and datasets (C) and heart samples coloured by tissues and (D) datasets.

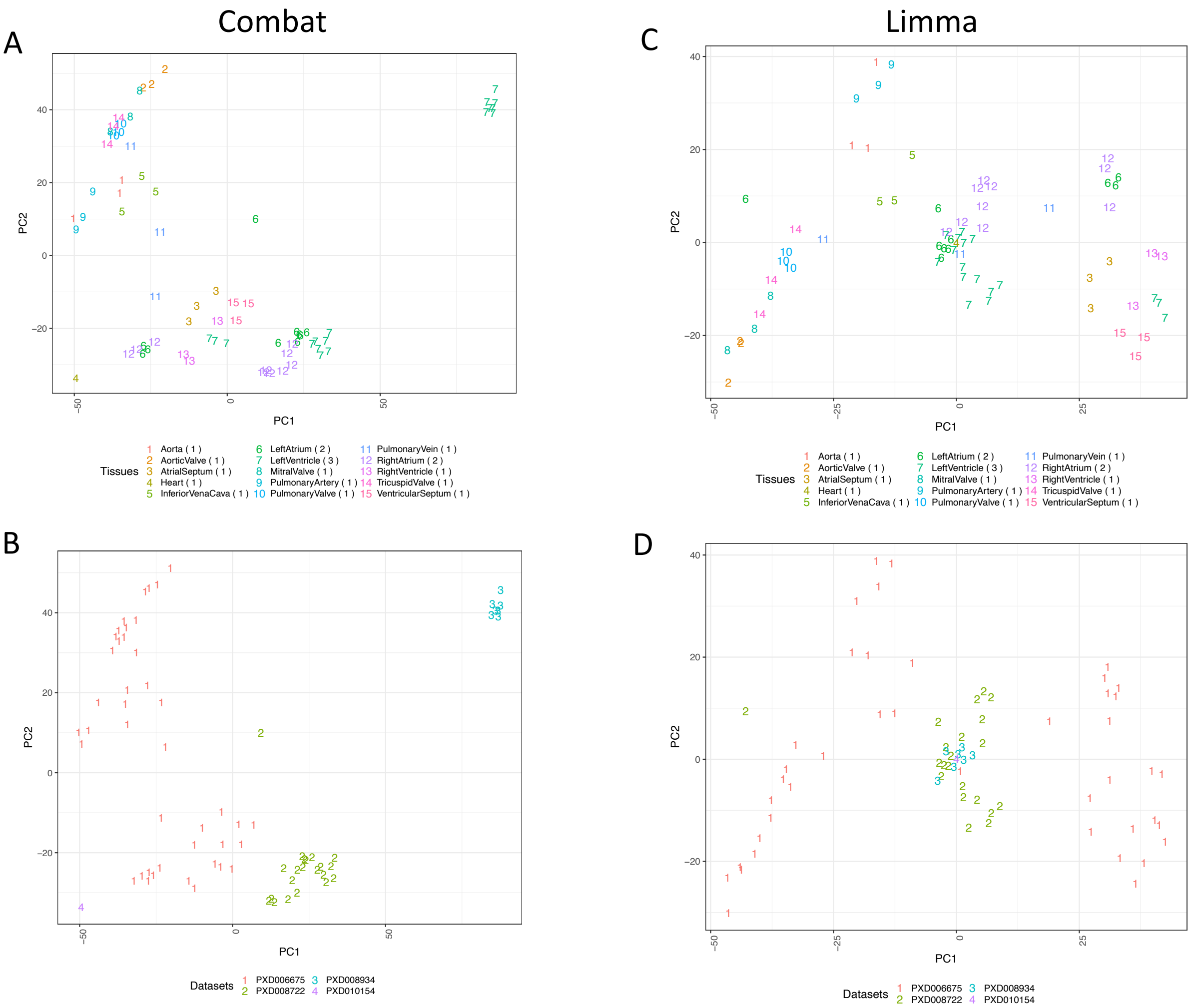


**Figure S3.** PCA of the heart samples using Combat normalisation where samples are coloured by (A) tissues and (B) datasets. Similarly PCA of heart samples using Limma normalisation where samples are coloured by (C) tissues and (D) datasets.

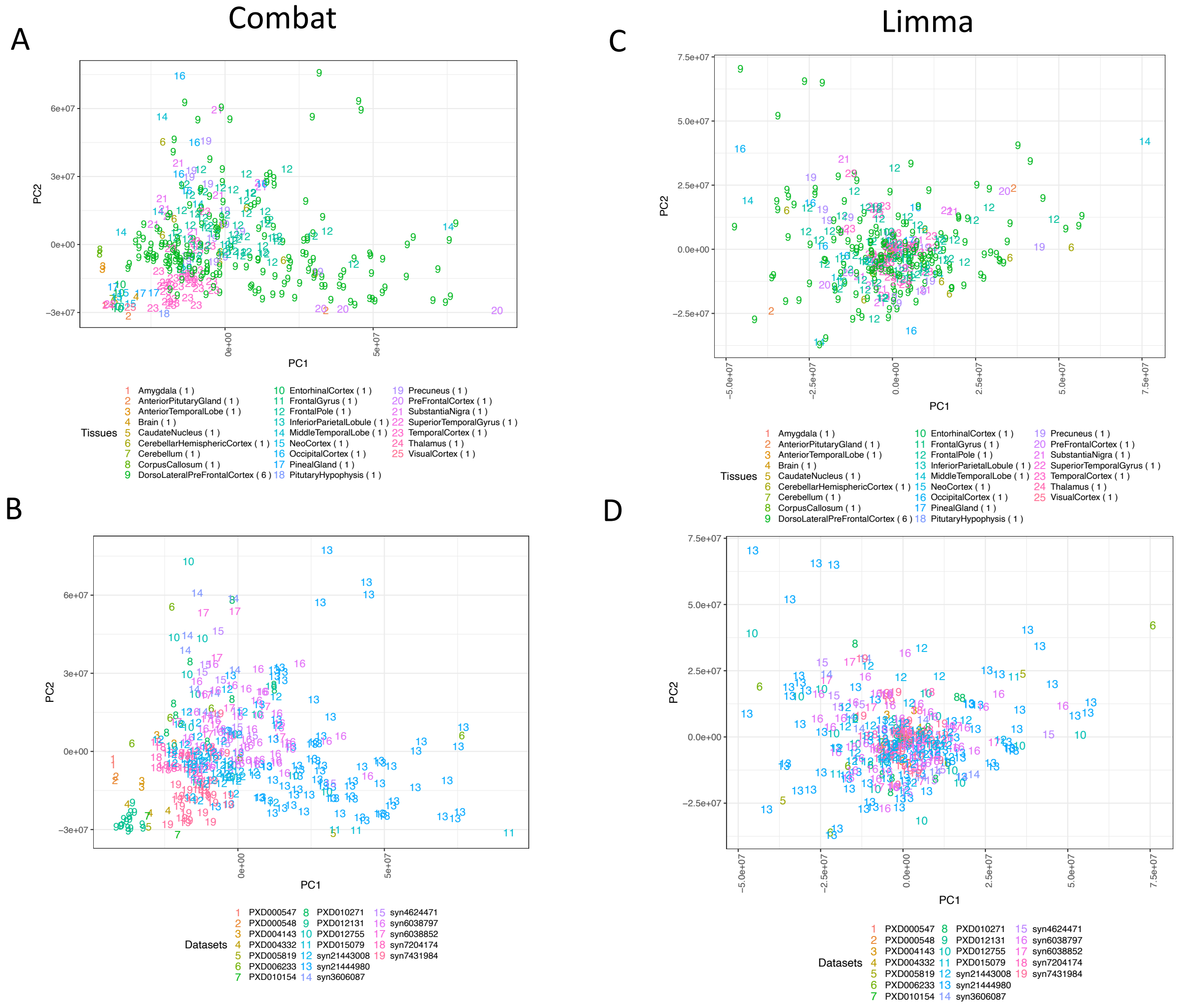


**Figure S4**. PCA of brain samples using Combat normalisation where samples are coloured by (A) tissues and (B) datasets Similarly PCA of brain samples using Limma normalisation where samples are coloured by (C) tissues and (D) datasets.

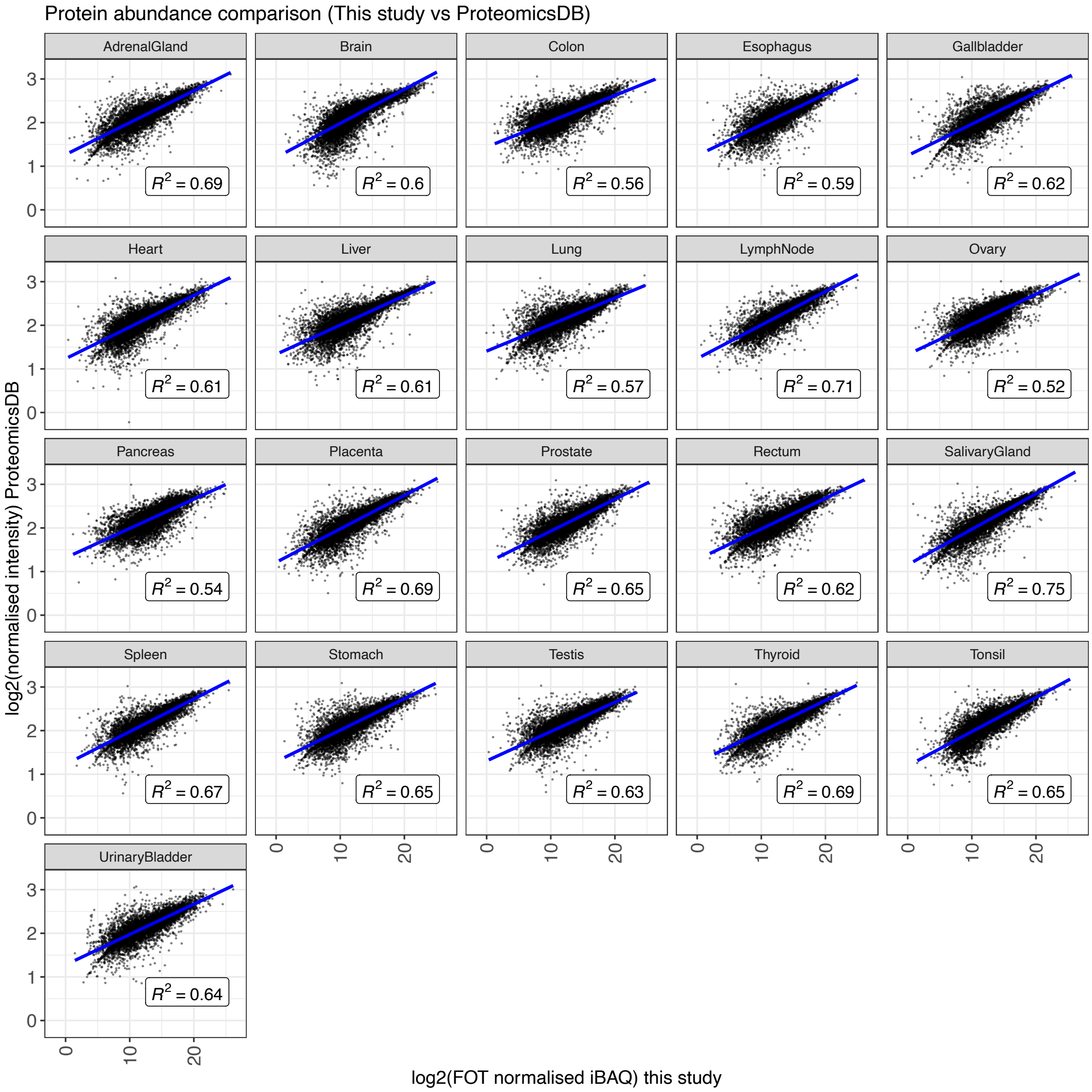


**Figure S5.** Correlation of protein abundances across various organs compared between ProteomicsDB and this study.

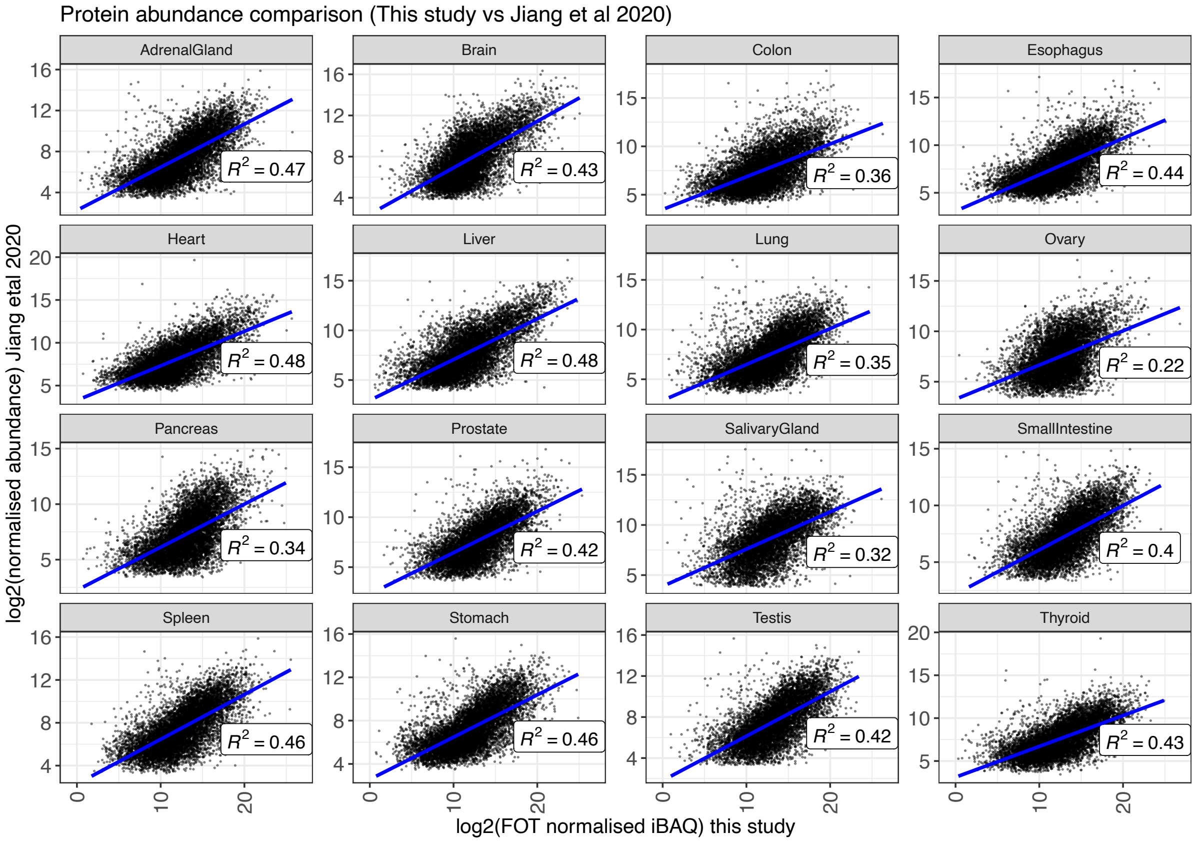


**Figure S6.** Comparison of protein expression between Jiang et al. (2020) (using TMT labelling method) and data presented in this study (label-free method).

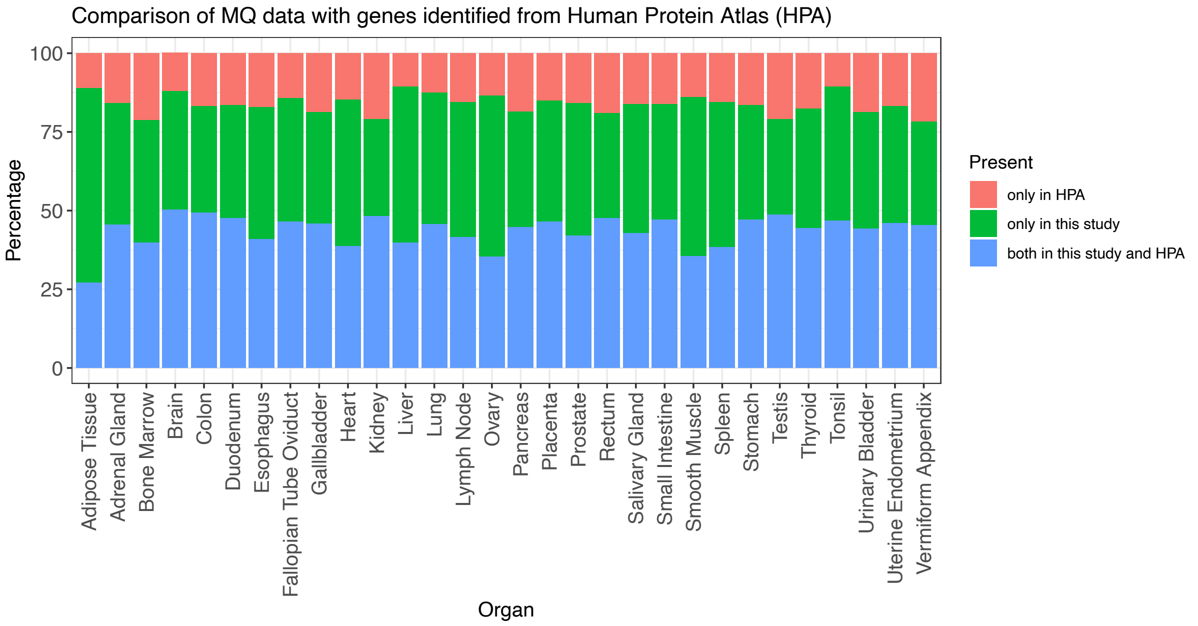


**Figure S7.** Protein detection comparison between Human Protein Atlas and the results included in this manuscript.

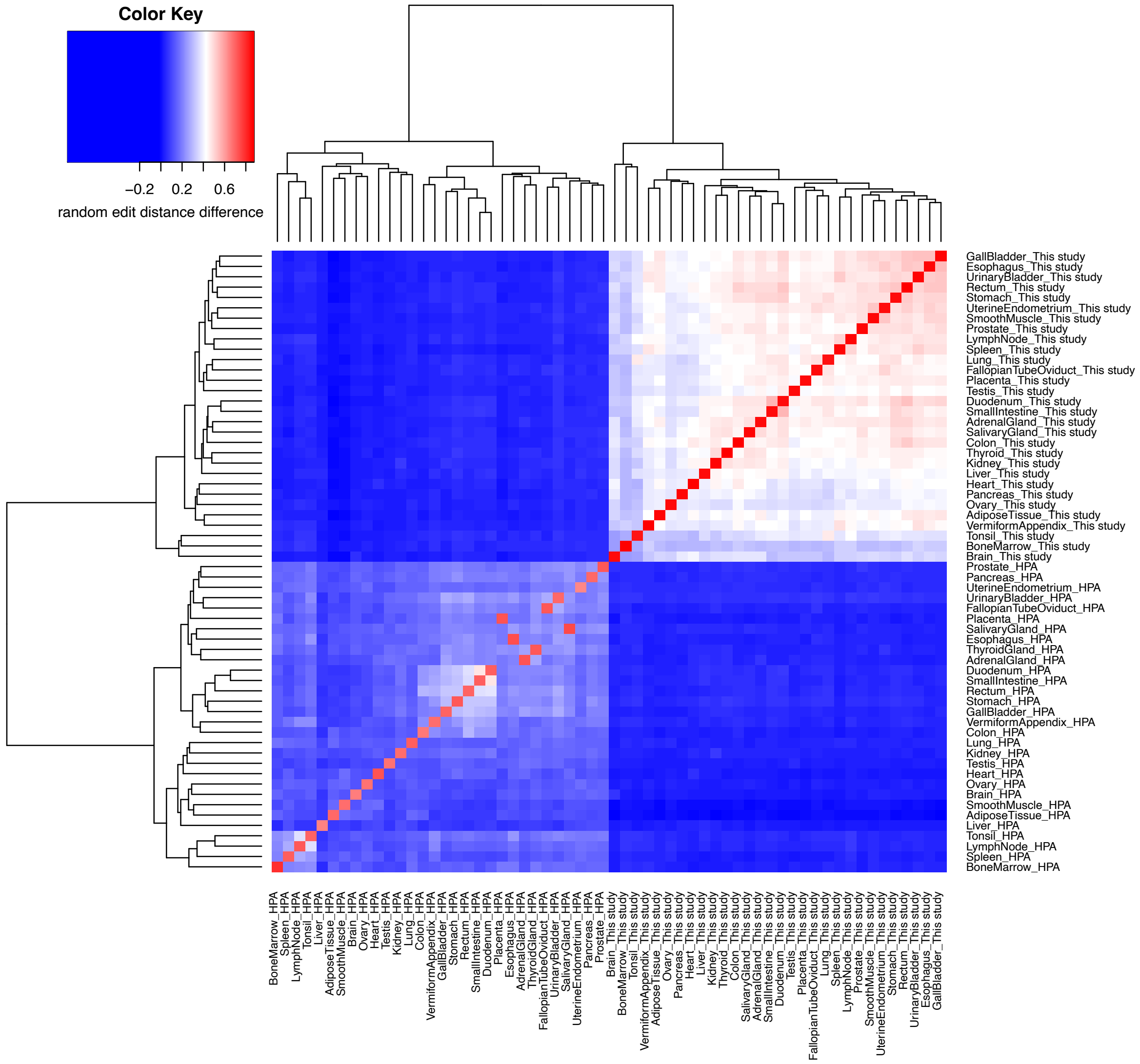


**Figure S8.** The randomised edit distance difference matrix showing the differences in the edit distances calculated for various organs between this study and the data from Human Protein Atlas.
